# Global disruption of plant biogeography by non-native species

**DOI:** 10.1101/2025.10.29.685360

**Authors:** Lirong Cai, Patrick Weigelt, Holger Kreft, Helge Bruelheide, Amy JS Davis, Wayne Dawson, Franz Essl, Mark van Kleunen, Ingolf Kühn, Bernd Lenzner, Jan Pergl, Petr Pyšek, Pieter B. Pelser, Jan J. Wieringa, Marten Winter

**Affiliations:** German Centre for Integrative Biodiversity Research (iDiv) Halle-Jena-Leipzig, 04103 Leipzig, Germany; Leipzig University, 04109 Leipzig, Germany; Department of Environmental Science, Radboud Institute for Biological and Environmental Sciences (RIBES), Radboud University, Heyendaalseweg 135, 6525AJ Nijmegen, The Netherlands; Biodiversity, Macroecology and Biogeography, University of Göttingen, 37077 Göttingen, Germany; Centre of Biodiversity and Sustainable Land Use, University of Göttingen, 37077 Göttingen, Germany; Campus-Institute Data Science, 37077 Göttingen, Germany; Institute of Biology / Geobotany & Botanical Garden, Martin Luther University Halle-Wittenberg, 06108 Halle, Germany; Ecology, Department of Biology, University of Konstanz, 78464 Konstanz, Germany; Department of Evolution, Ecology and Behaviour, Institute of Infection, Veterinary and Ecological Sciences, University of Liverpool, Liverpool, UK; Division of BioInvasions, Global Change & Macroecology, University Vienna, 1030 Vienna, Austria; Zhejiang Key laboratory for Restoration of Damaged Coastal Ecosystems & Zhejiang Provincial Key Laboratory of Plant Evolutionary Ecology and Conservation, Taizhou University, Taizhou 318000, China; Dept. Community Ecology, Helmholtz Centre for Environmental Research - UFZ, 06120 Halle, Germany; Department of Invasion Ecology, Institute of Botany, Czech Academy of Sciences, 25243 Průhonice, Czech Republic; Department of Ecology, Faculty of Science, Charles University, 12844 Prague, Czech Republic; School of Biological Sciences, University of Canterbury, Christchurch, New Zealand; Naturalis Biodiversity Center, Leiden, The Netherlands

**Keywords:** biogeographical regionalization, plant introduction, seed plants, dispersal limitation, environmental filtering

## Abstract

Biogeographical regions reflect differences in biotic composition resulting from long-term isolation and biogeographical processes, but how human-mediated introductions of non-native species have altered these biogeographical patterns remains unclear. Using global distribution data of 279,441 native and 10,067 non-native seed plant species, we analyzed the impact of species introductions on global organization of biogeographical regions. We show that species introductions have disrupted plant biogeography, causing the loss of distinct floristic realms and subrealms. Due to the impact of non-native species, geographic proximity and dispersal barriers are less important— particularly as trade-facilitated species exchange drives floristic homogenization—while environmental factors remain critical in structuring floristic regions. Our findings reveal that plant introductions erode biogeographical distinctness and highlight the urgent need for coordinated action to protect native biotas.

## Introduction

Biogeographical regionalization is a hierarchical system that categorizes geographic areas based on similarities in biotic composition, such that species or phylogenetic assemblages within each region are more similar to each other than to those in other regions (Holt *et al*., 2013; Morrone, 2018). Biogeographical regionalization thus provides a fundamental framework for addressing key questions in ecology, evolutionary biology, and conservation (Kreft & Jetz, 2010; Morrone, 2018). In plants, floristic regionalization was initially defined primarily by patterns of endemism (Takhtajan, 1986), recognizing six major floristic realms (Fig. S1), and later refined through the integration of species distribution data, phylogenetic information, and modern statistical approaches (Carta *et al*., 2022; Liu *et al*., 2023).

Biogeographical regions have resulted from millions of years of interplay of ecological, geological, and evolutionary processes. Two major processes have been shown to drive species distribution at large scales, thereby promoting the assembly of distinct biogeographical regions: (1) environmental filtering by physical, climatic and other abiotic factors, which determines where species can persist, and (2) dispersal limitations, which restrict species to their area of origin by preventing them from dispersing to environmentally suitable areas elsewhere, often reinforced by geographic barriers and geological processes such as plate tectonics (Liu *et al*., 2023; Cai *et al*., 2025). Any changes to these evolutionary and ecological processes would be expected to alter the world’s major biogeographical regions, influencing the future of plant diversity and the species that depend on it. Quantifying drivers and impacts of anthropogenic change to global biogeographical regionalization is therefore essential to anticipate, manage, and mitigate their consequences (Brown *et al*., 2023).

Species introductions are a major threat to biodiversity and drivers of global change (Pyšek *et al*., 2020). Approximately 16,500 vascular plant species (Davis *et al*., 2025), representing ∼ 4% of the global flora (Schellenberger Costa *et al*., 2023), are established outside their native range. Most plant introductions are linked to the exchange of traded goods, which enable species and their propagules to cross geographic barriers between historically isolated regions and result in the intentional (horticulture, breeding) or unintentional (contaminants, stowaways) introduction at unprecedented rates (Seebens *et al*., 2022). These plant introductions alter species composition and can lead to biotic homogenization, eroding the distinctiveness of regional floras (Winter *et al*., 2009; Daru *et al*., 2021; Yang *et al*., 2021). However, even after centuries of extensive human-mediated species exchange, the effects of plant introductions on the organization of global floristic biogeographical regions and the underlying mechanisms maintaining regional distinctiveness remain largely unknown, representing a critical gap for understanding and conserving Earth’s biodiversity.

Here, we provide the first comprehensive global assessment of the effects of plant introductions on natural plant biogeography and highlight the loss of distinct biogeographical regions. We tested for differences in the extent to which environmental filtering, dispersal limitation, and trade shaped plant regional compositions across the globe before and after plant introductions. Our study is based on a global dataset of regional species compositions for seed plants across 549 geographic regions worldwide (van Kleunen *et al*., 2019; Weigelt *et al*., 2020; Govaerts *et al*., 2021). After excluding ∼ 45,000 species from 293 genera that contain apomictic species, the datasets comprised 133,464 regional occurrence records for 10,067 naturalized non-native seed plant species that are establishing self-sustaining populations outside their native ranges, and 1,272,442 occurrence records for 279,441 native species (Fig. S2). By integrating species distribution data with a comprehensive megaphylogeny of seed plants (Smith & Brown, 2018), we quantified pairwise compositional dissimilarity between regions as taxonomic and phylogenetic turnover—the component of beta diversity independent of species or phylogenetic richness (Baselga, 2010)—to explore how the global spread of non-native plants has altered the natural biogeographical structure of the world’s flora.

## Results and Discussion

### Changes in global biogeographical regions

The spread of non-native species has significantly disrupted global natural biogeographical patterns of plants. Using a hierarchical clustering analysis of taxonomic and phylogenetic turnover, we identified highly distinct clusters of geographic regions as floristic realms and finer-scale clusters as subrealms, then compared floristic patterns before plant introductions (based on native seed plant occurrences only) and after introductions (including both native and non-native species; see Materials and Methods for more details). Our results show that before human-mediated introductions, the world’s flora was organized into 17 subrealms nested within seven realms based on taxonomic turnover (Fig. 1a-c), and into eight realms and 18 subrealms based on phylogenetic turnover (Fig. 2a-c). In contrast to Takhtajan’s six-realm floristic regionalization (Fig. S1) (Takhtajan, 1986), our delineation of natural floristic realms reveals clear boundaries between the Nearctic and Palaearctic realms, as well as between the African and Indo-Malesian realms, while other boundaries are mostly similar. Our delineation also largely confirms Liu et al.’s phylogenetically informed regionalization based on 12,664 angiosperm genera (Liu *et al*., 2023), although our phylogenetic scheme highlights the Cape as a distinct realm (Fig. 2b). Furthermore, we found that Madagascar clustered with the Indo-Malesian rather than the African realm, likely reflecting deep-time overseas dispersal of plant species following the separation of Madagascar from India 84–91 Ma and subsequent in situ diversification (Warren *et al*., 2010; Antonelli *et al*., 2022).

**Fig. 1.**
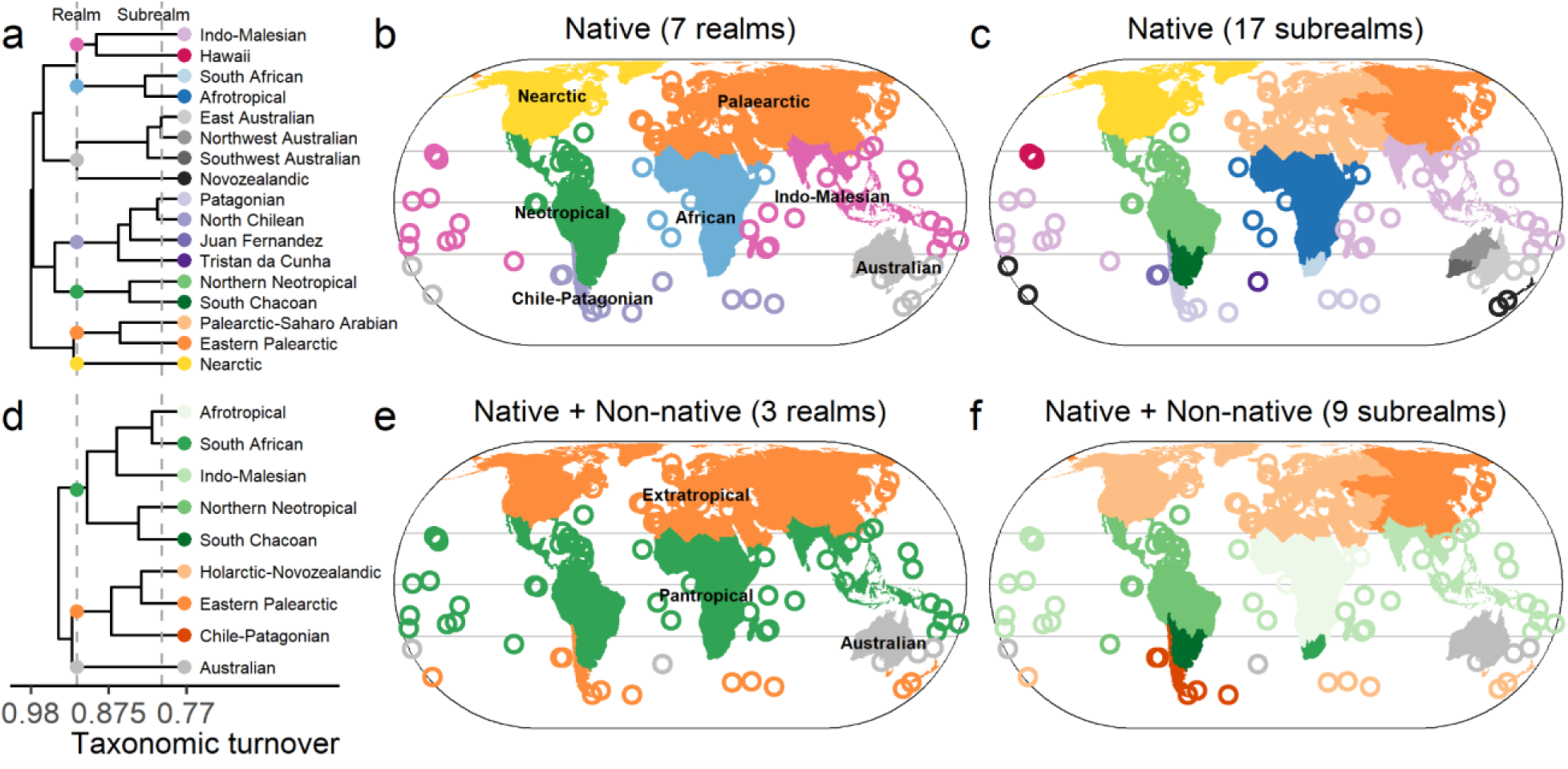
Global biogeographical patterns before and after plant introductions based on taxonomic composition. Taxonomic compositional dissimilarity was calculated using two species datasets: native species only (a–c) and all species (i.e., native and non-native species) combined (d–f), serving as the basis for hierarchical clustering. Dendrograms from hierarchical clustering (a, d) were pruned at two different heights to define biogeographical realms and subrealms. Realms are shown as distinct colors on the maps (b, e), while subrealms belonging to the same realm are depicted using consistent color gradients from the same palette on the maps (c, f). Matching colors are used to indicate corresponding realms and subrealms across the dendrograms and maps. Islands <50,000 km^2^ are represented by circles.

**Fig. 2.**
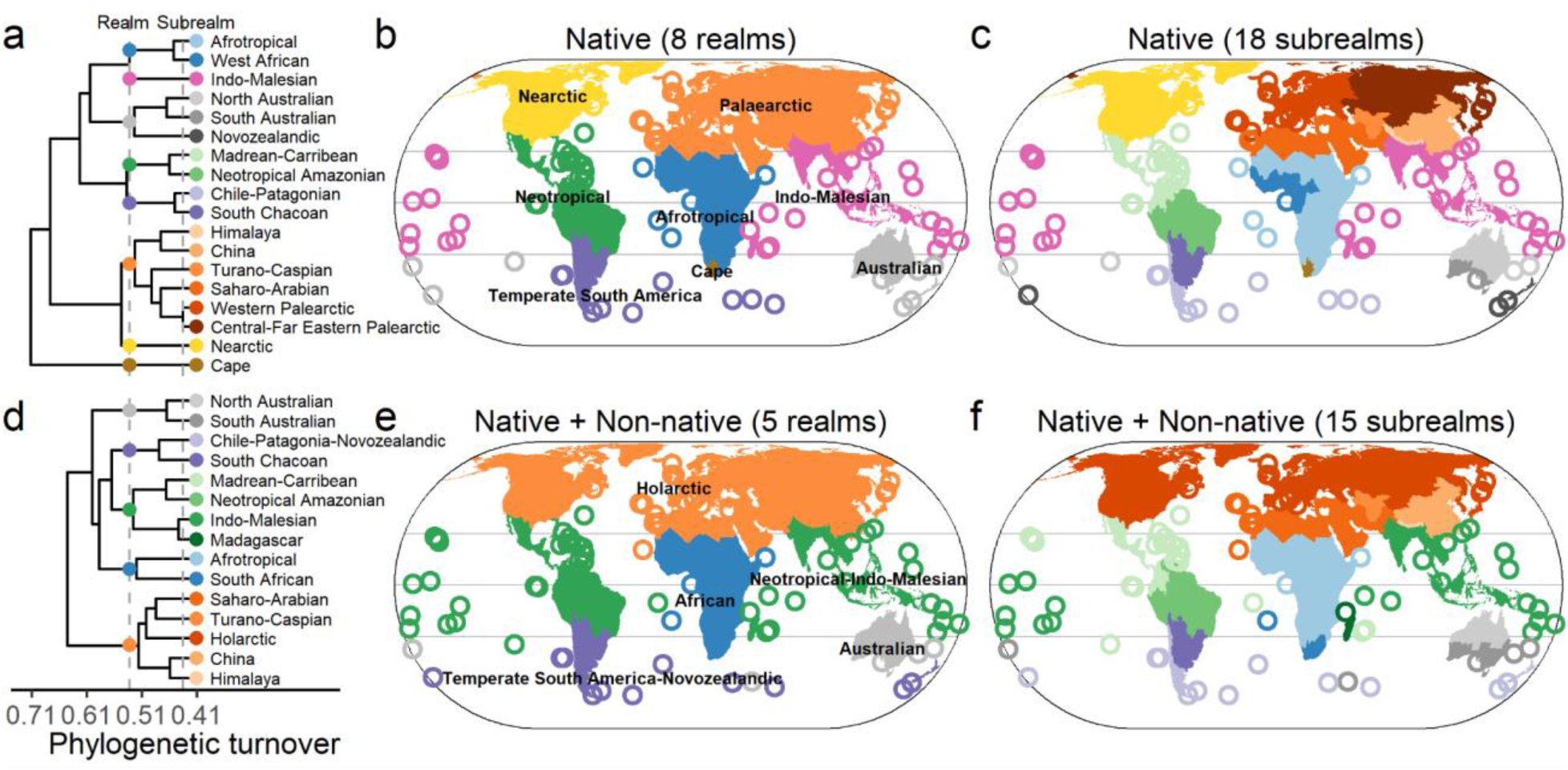
Global biogeographical patterns before and after plant introduction based on phylogenetic composition. Phylogenetic compositional dissimilarity was calculated using two species datasets: native species only (a–c) and all species (i.e., native and non-native species) combined (d–f), serving as the basis for hierarchical clustering. Dendrograms from hierarchical clustering (a, d) were pruned at two different heights to define biogeographical realms and subrealms. Realms are shown as distinct colors on the maps (b, e), while subrealms belonging to the same realm are depicted using consistent color gradients from the same palette on the maps (c, f). Matching colors are used to indicate corresponding realms and subrealms across the dendrograms and maps. Islands <50,000 km^2^ are represented by circles.

Due to human-mediated plant introductions, however, this regionalization collapsed: four realms and eight subrealms disappeared from the taxonomic regionalization when the dissimilarity threshold used to define floristic regions was held constant (Fig. 1d-f). Tropical and subtropical regions spanning America, Africa, and Asia now share a higher proportion of species and were grouped into a single Pantropical realm (Fig. 1e). Also, temperate to Arctic areas in both hemispheres, along with surrounding islands, coalesced into an Extratropical realm, except for Australia, which emerged as a separate realm following the exclusion of the Novozealandic subrealm. Consequently, our regionalization showed that plant introductions restructured the global flora into three biogeographical realms: a (sub)tropical belt (Pantropical), a unified temperate-Arctic region (Extratropical), and the Australian realm.

The loss of formerly distinct subrealms was particularly pronounced for islands (Fig. 1f). Islands are renowned for their high levels of endemism and floristic differentiation (Schrader *et al*., 2024), shaped by dispersal filtering, in-situ speciation, and lineage persistence over long timescales due to isolation (Gillespie & Roderick, 2002). Our findings indicate that the introduction of non-native plant species has diminished the distinctiveness of floras on some remote islands (Castro *et al*., 2010). Notably, islands or archipelagos with an exceptional level of endemism (Schrader *et al*., 2024) that were previously classified as distinct subrealms—such as Hawaii and the Juan Fernández Islands—now show increased similarity in species composition with nearby mainland areas or other islands (Fig. 1f). As a result, these islands have been reclassified and merged with adjacent regions. In contrast, New Zealand merged with the geographically distant Nearctic and Palearctic–Saharo-Arabian subrealms, reflecting species translocations linked to Europe’s colonial history (Lenzner *et al*., 2022). Such breakdowns in biogeographical distinctiveness were also observed for some mainland regions, such as the Southwest Australian subrealm, which, despite its unique geological history and high endemism (Crisp *et al*., 2001; Cai *et al*., 2023b), also experienced a loss of floristic differentiation and the breakdown of its natural boundaries with other pan-Australian subrealms. Among non-native taxa, certain species became established across most or all areas within the colonized realms, thereby strongly contributing to blurring of inter-realm distinctions (Table S1). For example, *Rumex acetosella*, a Eurasian plant in the Polygonaceae family, was introduced across the Juan Fernández and Patagonian subrealm and contributes to their floristic homogenization (Fig. S3).

When accounting for evolutionary relationships, we also found a loss of realms and subrealms (though less pronounced than for the taxonomic level) from eight to five and 18 to 15, respectively (Fig. 2). Transoceanic plant introductions facilitated the emergence of novel transoceanic realms and subrealms, such as the Indo-Malesian uniting with the Neotropical realm, the Nearctic uniting with the Palaearctic realm, as well as the Novozealandic subrealm uniting with the Chile-Patagonian realm (Fig. 2e and f). Our comparative analyses revealed that plant introductions have resulted in more pronounced losses of formerly distinct floristic regions at the taxonomic than at the phylogenetic level. This discrepancy is likely due to the broader distribution of certain seed plant lineages at higher taxonomic level (e.g., families) that are less constrained by dispersal limitation because they had more time to spread worldwide over evolutionary timescales and could occupy large parts of their potential range, resulting in higher phylogenetic similarities among regions that harbor different species within the same lineage (Cai *et al*., 2025). In contrast, the taxonomic approach emphasizes species-level differentiation, which is more sensitive to compositional differences irrespective of the degree of relatedness among those species. For natural patterns, the Cape emerged as a distinct realm only based on phylogenetic composition (Fig. 1b and Fig. 2b), reflecting its evolutionary uniqueness with over 160 endemic genera, many of which are highly species-rich due to an extensive history of species radiations (Goldblatt *et al*., 2005; Linder, 2005).

To assess the influence of potentially incomplete non-native species data used in this study, particularly in tropical regions, we performed sensitivity analyses by randomly adding a fixed number of non-native species from the distribution dataset to checklists of likely undersampled regions, repeating the taxonomic composition analysis 1,000 times (see Materials and Methods for more details). Our results showed that adding non-native species largely preserved the original patterns (i.e., three realms, nine subrealms) but occasionally intensified floristic homogenization, reducing the global realms to two: a (sub)tropical belt (Pantropical) and a unified temperate-to-arctic region (Extratropical) (Fig. S4), as found for gastropods (Capinha *et al*., 2015). Given the likelihood of under-documented non-native floras in some tropical regions (Chong *et al*., 2021), our results may underestimate the real extent to which non-native species have already eroded natural biogeographical regions. As more non-native species are recorded, their role in reshaping global biogeographical patterns is likely to become even more pronounced. We also likely underestimate the disruption of global plant biogeography because plant extinctions were not considered in our study, as extinction data are unavailable for many regions. Although extinct taxa are fewer in number per region and thus exert limited effects on overall patterns, most had narrow ranges, contributing to the loss of regional floristic distinctiveness (Winter *et al*., 2009). Further, to assess the robustness of the pattern we found and to avoid dominating effects of well-documented regions, we performed 1,000 iterations of the taxonomic composition analysis, each time randomly removing non-native species from regional checklists with more than 80 non-native plant species. We consistently observed a reduction in the number of realms and subrealms due to plant introductions, with a high frequency in the clustering results of four realms and 10 subrealms (Fig. S5). This indicates that despite regional differences in the completeness of non-native species records, their disruptive effects on biogeographical patterns remained detectable, confirming that the detected loss of formerly distinct floristic regions was robust.

### Changes in drivers of compositional dissimilarity due to plant introduction

Our analysis demonstrates that plant introductions disrupt the natural and fundamental biogeographical processes, such as dispersal limitations, governing plant distributions across the globe. We assessed the relative contributions of environmental filtering and dispersal limitation in shaping compositional dissimilarity before and after plant introductions by modeling turnover of three species groups (i.e., native species only, non-native species only, and all species combined) as a function of environmental dissimilarity and geographical distance (i.e., geographical linear distances and least-cost distances accounting for geographic barriers) using generalized dissimilarity models (Ferrier *et al*., 2007) (Fig. 3; Figs. S6 and S7). Deviance partitioning of the models (Fig. 3a) revealed that the contribution of geographical distance to taxonomic and phylogenetic turnover markedly decreased when the results for the native-only species group (43.1% and 23.2% of the deviance for taxonomic and phylogenetic turnover, respectively) are compared with those for the non-native (5.4% and 1.7%) and all-species groups (22.8% and 15.6%). This decline indicates that human-mediated plant introductions have weakened natural dispersal limitations, diminishing the role of geographical distance in structuring global plant distributions.

**Fig. 3.**
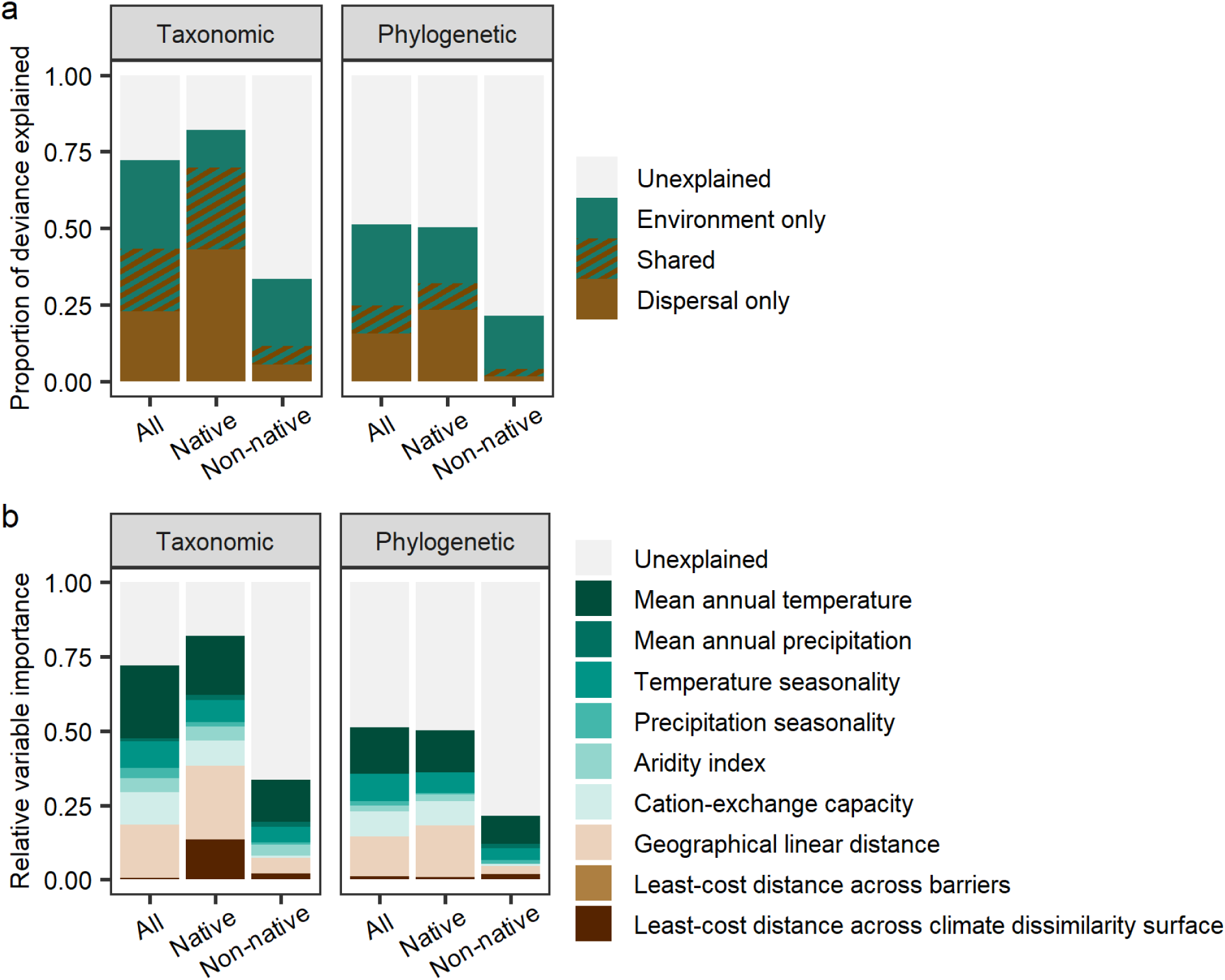
Relative importance of predictor variables for plant taxonomic and phylogenetic turnover in seed plants. In (a), relative importance is shown for dispersal and environment-related groups of predictor variables based on deviance partitioning for all, native, and non-native species. In (b), relative importance is shown for individual predictor variables based on the height of generalized dissimilarity modeling transformation curves, which is scaled so that their sums equal the proportion of deviance explained by the model.

Specifically, the effect of geographical linear distances, a proxy for geographic proximity, was markedly reduced, from 24.7% for natives to 5.1–17.9% when non-native species were included in taxonomic turnover, and from 17.4% to 2.8–13.3% in phylogenetic turnover (Fig. 3b and Fig. S6). However, geographical linear distances do not necessarily capture all constraints on species dispersal imposed by landscape features. To address this, we explored alternative distance measures that take into account physical and environmental barriers, including water, mountains, deserts, or unsuitable climates, along a hypothetical continuous dispersal path between two regions (Cai *et al*., 2025). We found that least-cost distances across climatically dissimilar surfaces, which quantify the barrier effect of climatically unsuitable habitats between regions, strongly influenced taxonomic turnover for native species (13.5%; Fig. 3b). However, this effect declined sharply when non-native species were included (0.5 and 2.1%; Fig. 3b). Therefore, our findings demonstrate that plant introductions enable movements of species between regions that are geographically distant and separated by climatic and physiogeographical barriers, particularly where natural corridors or climate-similar stepping stones are absent, leading to the disruption of plant biogeography. These effects are especially disruptive on islands, where dispersal has historically been restricted by oceans that offer no viable pathways for plant colonization, yet has been overcome through human-mediated introductions (Helmus *et al*., 2014). Overall, our results on the effect of geographical distance on compositional similarities show that biological introductions diminish the effects posited by the classical biogeographical law of distance decay (Nekola & White, 1999; Capinha *et al*., 2015) and reduce the isolating effects of geographic barriers along dispersal routes.

In contrast, environmental dissimilarity consistently exerted a strong positive influence on both taxonomic (12.3-28.9%) and phylogenetic turnover (17.3-26.4%) across all three species groups, and even increased for all-species groups (taxonomic turnover: 28.9%; phylogenetic turnover: 26.4%) compared to native-species (12.3%; 18.2%; Fig. 3a). Among environmental variables, climatic factors—especially annual mean temperature (14.1-24.6%; 9.4-15.8%)—emerged as the most important drivers of compositional dissimilarity (Fig. 3b). These findings indicate that although human activities enabled species to overcome natural dispersal limitation, the successful establishment and persistence of non-native species remain strongly constrained by environmental, especially climatic conditions of the colonized region. Such climate-driven filtering partially explains why a clear boundary between the Pantropical and Extratropical realms exists when the distribution patterns of both native and non-native species are considered (Fig. 1d and e). However, ongoing global climate change is likely to alter these environmental constraints, potentially accelerating shifts in biogeographical patterns—driving poleward movement of biogeographical boundaries, reducing the extent of extratropical realms, and amplifying the homogenizing effects of plant introductions (Minev-Benzecry & Daru, 2024).

### The effects of trade flow on floristic distinctiveness changes

To directly assess the role of anthropogenic dispersal in driving biogeographical change, we calculated bilateral trade flows between pairs of regions (Gaulier & Zignago, 2010) and evaluated their effects on regional compositional dissimilarity change. Although the majority of plant introductions occurred in the last 250 years (IPBES, 2023) and the best available trade data we used are relatively recent, it has been shown that present-day trade connections and their relative sizes within the global trade network likely mirror historical patterns of economic exchange—varying locally but being consistent globally (Mitchener & Weidenmier, 2008; Foti *et al*., 2013; Gokmen *et al*., 2020). When modelling the relative contribution of bilateral trade flows on turnover, we found a weak but detectable effect for non-native and all-species groups (Figs. S8 and S9). However, this result should be interpreted with caution, as trade data were available for only 8,417 of the 115,921 region pairs and then we rescaled data to subregions and assign zeros to missing values for modelling, which may have introduced bias (see Materials and Methods for more details).

To further assess the role of trade flow, we quantified the difference in the number of shared species between regions after introductions, calculated as post-introduction minus pre-introduction values, and modeled it against bilateral trade flows and climatic distance for region pairs with trade data. We found that bilateral trade flow emerged as the stronger predictor, significantly increasing shared species richness among regions (Fig. 4). These results indicate that stronger trade connections substantially enhance species exchange between regions, facilitating the reshaping of global plant biogeography. For instance, the Juan Fernández Islands—once an isolated and uninhabited archipelago—became integrated into global trade networks after Spanish discovery and subsequent integration into Chile, leading to a continuous influx of goods, people, and introduced plants that ultimately contributed to the breakdown of the formerly distinct Juan Fernández and Patagonian subrealm. Further, climatic distance had a significant negative effect on shared species richness (Fig. 4), indicating the joint effects of human-mediated dispersal and environmental suitability in shaping global plant introductions.

**Fig. 4.**
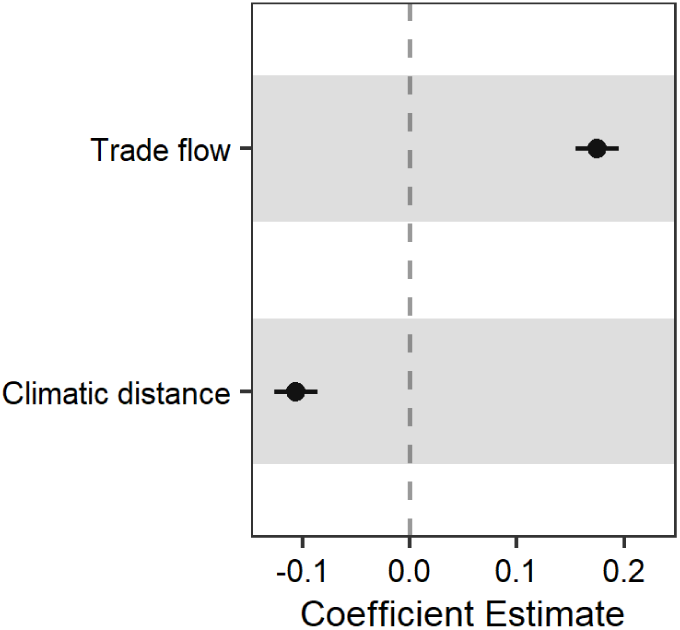
Determinants of difference in species numbers shared by pairs of regions after plant introductions based on regression models. Difference in shared species is calculated as the number of shared species after introduction minus that before introduction. Shown are standardized regression coefficients for each predictor variable, with bars around each point indicating standard errors of the coefficient estimate. Each predictor variable was standardized to zero mean and unit variance to aid model fitting and make their parameter estimates comparable.

## Conclusion

Here, we demonstrate how introductions of new plant species by humans have disrupted millions of years of biogeographical structuring of the global flora, breaking down long-standing natural barriers and eroding biogeographical distinctness of regional species pools. These breakdowns are very likely to continue and accelerate. With ongoing globalization and its associated global trade, monitoring the flux of species over natural barriers is a daunting global challenge, intensified by incomplete introduction knowledge in many countries (McGeoch *et al*., 2023; Davis *et al*., 2025). The conservation-relevant management of newly arriving species and their diverse introduction pathways across the planet is an even bigger task. The global research and decision-making communities need to increase their efforts to monitor and manage highly dynamic species compositions and major introduction pathways for plants, by prioritizing regions of high floristic uniqueness such as islands with high endemism and tropical areas currently with low numbers of non-native species. Over the next decade, coordinated investments in global species monitoring, interoperable data systems, and cross-border management of biological introductions will be essential to mitigate the ongoing breakdown of the world’s natural floristic biogeography.

## Materials and Methods

### Species distribution data

We compiled a comprehensive global dataset on the regional species composition of native and naturalized non-native seed plants by integrating three major sources. For native species, we used the Global Inventory of Floras and Traits (GIFT version 3.0: http://gift.uni-goettingen.de) (Weigelt *et al*., 2020) and the World Checklist of Vascular Plants (WCVP: https://powo.science.kew.or) (Govaerts *et al*., 2021). Both databases provide comprehensive vascular plant distribution data, derived from published floras, checklists, and online databases. GIFT provides plant species inventories for ∼3,400 geographic regions worldwide, including islands, protected areas, biogeographical regions, and political units (e.g., countries or states). WCVP provides taxonomically standardized data on accepted vascular plant species and their distributions across 369 botanical countries. For naturalized non-native species, we used the Global Naturalized Alien Flora (GloNAF version 2.02: https://zenodo.org/records/14696776) (van Kleunen *et al*., 2019), which includes checklists of naturalized non-native species in vascular plants for ∼1,300 geographic regions (e.g., countries, islands) globally. Naturalized non-native species were defined as those establishing self-sustaining populations outside their native ranges (Richardson *et al*., 2000). Geographic regions in GIFT and WCVP provided complementary coverage, and when combined, offered a globally comprehensive dataset of native plants, which allowed for complete matching with the regional setting of GloNAF. In addition to GloNAF, we used distribution information from WCVP for non-native plant species as a complementary dataset to fill gaps in the global coverage of non-native species distribution datasets (Fig. S2d). Regional inventory data were preferred here because they offer the most complete and authoritative records of regional floristic composition, whereas point occurrence datasets (e.g., GIBF) tend to be geographically biased and cover only a fraction of regional floras (Meyer *et al*., 2016; Daru & Rodriguez, 2023). Because all nonhybrid species names in GIFT 3.0 and GloNAF 2.02 were standardized and validated based on taxonomic information provided by WCVP, we were able to directly combine the three datasets. Although the WCVP dataset may contain records of casual non-native species in addition to naturalized non-native species, we opted to include these data to address data gaps present in the GloNAF database (Fig. S2c and d), given that the number of non-naturalized species is likely relatively low in the WCVP dataset. Moreover, a large fraction of casual species are highly likely to become naturalized, which would reinforce the main patterns we found.

To obtain a dataset consisting of regions with distribution information for both native and non-native species, we conducted the following procedures. First, we downloaded native seed plant distribution data from both GIFT (Denelle *et al*., 2023) and WCVP (accessed 18 February 2022), and merged the records of species distribution from GIFT with those from WCVP available for the same regions. We removed the regions with highly incomplete native seed plant checklists which were assessed by the GIFT curators. Next, we identified regions with a complete spatial match between GloNAF and the native species dataset (GIFT/WCVP). For regions that did not align perfectly, we merged smaller GloNAF regions, which were nested within larger regions in the native species dataset and fully covered them, to match the regions in the native species dataset, and vice versa. In these merged regions, species were classified as native if they were native in any of the smaller nested regions. Conversely, species were considered non-native in the merged region if they were non-native in any of the smaller regions and not native in the larger merged region. To ensure high data quality and complete global spatial coverage, we selected and handled regions in three steps based on the completeness of GloNAF species checklists. First, we included only regions where at least one species list was judged to include more than 50% of the naturalized non-native taxa for that region in GloNAF. Second, for regions with less than 50% of the naturalized non-native species, we merged non-native species distribution data from WCVP with the records in GloNAF for the same regions. Third, we added regions that had non-native species distribution data only from WCVP (16 out of 549 regions) to fill gaps. Although the assessment of species-list completeness was mainly conducted by the GloNAF curators and might be coarse, it allows us to identify regions with a high likelihood of significant data deficiencies.

To resolve conflicts in species status across datasets, we applied the following data-cleaning procedures: (1) when two datasets agreed on a species’ status and the third differed, we retained the consensus status—for example, if a species was classified as native in both GIFT and WCVP but as non-native in GloNAF, we retained the native status; (2) when status differed between GIFT and WCVP or between WCVP and GloNAF, we prioritized native classifications in GIFT or WCVP and excluded corresponding non-native records from WCVP and GloNAF because WCVP provides additional comprehensive and very reliable distribution data for native species but not for non-native species (Govaerts *et al*., 2021); (3) when status differed between GIFT and GloNAF, we retained the non-native classification from GloNAF because we consider this the most reliable database for distribution data of non-native species, and excluded the opposing native plant record from GIFT. Before applying the status harmonization procedure, we refined the GIFT dataset by removing 15,018 native distribution records (out of 1,287,460) that fell outside the continental-level native range as defined by the WCVP dataset for each species. The final dataset included 324,436 seed plant species from 549 non-overlapping regions, including 438 mainland regions and 111 islands or island groups (Fig. S2f). The geographic regions in the dataset completely covered the globe, and varied in size from 17 to 3,069,766 km^2^ (median: 92,098 km^2^). To assess whether the revision of native distributions in GIFT using WCVP native range influenced the observed patterns, we additionally performed analyses using the dataset without the removal of native distribution records from GIFT (Figs. S10 and S11).

### Apomictic taxa

Apomixis is a form of asexual seed reproduction that occurs without gamete fusion. This reproductive strategy is often linked to hybridization and polyploidy, promoting reticulate evolution and the emergence of novel lineages (Majeský *et al*., 2017). The taxonomic treatment of apomictic taxa—especially species delimitation—remains debated, with inclusion and levels of taxonomic resolution varying across regional floras and checklists. This discrepancy can introduce geographical biases in the global distribution of apomictic taxa, especially towards the well-studied European floras (Richards, 2003), leading to greater unique compositions in regions with a high proportion of recognized apomictic taxa (Cai *et al*., 2025). To address the bias introduced by apomictic taxa, we excluded all species from 293 genera that contain apomictic species (44,995 species), as identified in the Apomixis Database (Hojsgaard *et al*., 2014). The resulting dataset, used for the main analyses, included 279,441 species. While the Apomixis Database focuses exclusively on apomixis in angiosperms, it is well-established that apomixis is very rare in gymnosperms (Majeský *et al*., 2017).

To compare biogeographical patterns accounting for non-native species, we categorized the species distribution data into three groups: all species, native species, and non-native species. The native species dataset contained 1,272,442 distribution records of 279,441 native species (i.e., those present in a region before human-mediated dispersal), the non-native species dataset included 133,464 records of 10,067 non-native species (i.e., present in the region as a result of dispersal mediated by humans), and the all-species dataset (including both native and non-native species) contained 1,405,906 records of 279,441 species (i.e., present as a result of both natural and human-mediated dispersal).

### Phylogeny

To assess phylogenetic turnover, we mapped species from the distribution dataset to the large, dated species-level phylogeny of seed plants (Smith & Brown, 2018), which encompasses 353,185 terminal taxa. Of the species in the distribution dataset, 215,278 species (77.0%) were already part of the original phylogeny. For the remaining species, we conservatively assigned them to their respective congeners within the phylogeny by replacing all species of a given genus with a polytomy, utilizing the ‘congeneric.merg’ function from the R package ‘pez’ (Pearse *et al*., 2015). We then excluded species not found in the distribution dataset from the phylogeny, creating a merged phylogeny that includes 272,511 out of 279,441 species from the distribution dataset (97.5%). Although adding polytomies may introduce additional phylogenetic uncertainties, it has been shown that there is a strong correlation between phylogenetic metrics based on phylogenetic trees with many polytomies and those based on trees with no or fewer polytomies (Qian & Jin, 2021). To assess whether incorporating species by replacing their genera with polytomies affects phylogenetic turnover patterns, we calculated phylogenetic turnover using a matched phylogeny that contained only 215,278 species present in the original phylogeny and without unplaced species added. The results revealed a strong correlation between phylogenetic turnover calculated from the merged and matched phylogenies (Pearson’s r = 0.99), demonstrating the robustness of the results based on the merged phylogeny. Consequently, the merged phylogeny was utilized for all analyses presented in the main text.

### Taxonomic and phylogenetic turnover

We used the Simpson dissimilarity index to measure variation in species and evolutionary composition between all pairs of regions (Baselga, 2010; Leprieur *et al*., 2012). This index captures the turnover component of beta diversity, defined as species or lineage replacement between regions (Baselga, 2010; Leprieur *et al*., 2012). This index is insensitive to variations in species and phylogenetic richness, allowing for the comparison of regions with differing area sizes that exhibit significant differences in species and phylogenetic richness (Cai *et al*., 2023a). The index is defined as: ß_sim_= min(b,c)/[min(b,c) + a]. When taxonomic turnover is calculated, a is the number of species shared by two regions, and b and c are the numbers of species unique to each region, respectively. For phylogenetic turnover, species numbers are replaced with total branch lengths in the phylogeny (Leprieur *et al*., 2012). In addition to turnover, we calculated the number of species shared by each pair of regions. All metrics were computed separately for the native, non-native, and all-species datasets.

### Potential drivers of taxonomic and phylogenetic turnover

To test how the effects of environmental filtering and dispersal limitation on taxonomic and phylogenetic turnover vary with and without non-native species, we identified a set of predictor variables representing environmental dissimilarity, geographical distances, and anthropogenic dispersal.

#### Environmental factors

To capture ecologically relevant dimensions of environmental space for seed plants, we considered climate and soil variables, which have been shown to be important to regional compositional dissimilarity of plants (Eiserhardt *et al*., 2013; Cai *et al*., 2025). We included five climatic variables of ecological relevance (Cai *et al*., 2025), namely mean annual temperature, mean annual precipitation, temperature seasonality, precipitation seasonality from CHELSA (Karger *et al*., 2017), and aridity index (i.e., the ratio of precipitation to potential evapotranspiration) from Global Aridity Index and Potential Evapotranspiration Database (Zomer *et al*., 2022). We also included soil cation-exchange capacity (cmol +/kg) as a proxy for soil fertility from SoilGrids250m (Hengl *et al*., 2017). For climatic and edaphic variables, we extracted the mean value of each variable for each region from the corresponding raster layers.

#### Geographic distances

To quantify the effects of dispersal limitation on turnover, we included geographical linear distances among regions, the least-cost distance across barriers, and the least-cost distance across a surface of climate dissimilarity (Cai *et al*., 2025). To calculate distances between pairs of regions that differ in area, we calculated centroid-to-centroid distances between each pair of grid cells, which are equal-area and equidistant hexagons (grid area: 23,323 km^2^) across the Earth’s surface (Barnes & Sahr, 2017). Pairwise distances between regions were then measured as the minimum distance between the grid cells overlapping with the first and second focal regions. We calculated geographical linear distances among regions as the shortest distance between each pair of grid cells on the WGS 84 reference ellipsoid (i.e., great-circle distance).

Following Cai et al. (2025), we calculated the least-cost distance across barriers and across climate dissimilarity surface for each pair of grid cells using network analysis. Least-cost distance between regions quantifies the minimum cost of moving from one region to another across cost surfaces. Network analysis can identify minimum-cost paths between vertices in a network, where vertices represent entities and edges represent the connections between the vertices and can be weighted. A path between two vertices consists of a sequence of edges from the first vertex to the second, and the total length of the path is the sum of the edge weights along that path (McNulty, 2022).

First, we calculated the least-cost distance across barriers considering water bodies, mountains and cold and dry deserts as dispersal barriers. We constructed a network for equal-area, equidistant hexagon grids using the igraph R package, with each grid cell connected to its six immediate neighbors. We calculated the weights of edges as geographical linear distances between two connected grid cells multiplied by the mean of the values extracted from a given barrier-cost surface for the two grid cells. We defined four barrier-cost surfaces based on water presence, mean elevation (Danielson & Gesch, 2011), annual mean temperature, and aridity index, to quantify the costs of crossing water bodies, mountains, and cold and dry deserts. Barrier costs were scaled from 0 to 1, with higher values indicating higher costs for moving from one grid cell to another. To account for multiple barriers simultaneously, we combined the four barrier-cost surfaces into a single barrier-cost surface by taking the maximum value across the four surfaces for each grid cell. Then, we calculated the least-cost distance between each pair of grid cells as the length of the shortest path between pairwise grid cells for the combined barrier-cost surface. Finally, least-cost distances between geographic regions were obtained as the minimum distance between all grid cell pairs overlapping the two regions.

We also calculated the least-cost distance across climate dissimilarity surface to quantify dispersal limitation due to climatically unsuitable habitats. Unlike the previous barrier costs, the climate cost surfaces were defined individually for each grid cell to capture the specific climatic conditions of the focal region. Each grid cell’s climate was characterized using the first five axes of a principal component analysis of 19 CHELSA bioclimatic variables (Karger *et al*., 2017), with precipitation-related variables log-transformed; these axes captured 93.5% of climate variability. For each focal grid cell, climate costs to all other grid cells were defined as the Euclidean distance of the five climatic axes between each grid cell and the focal grid cell, and scaled to [0,1] by dividing by the maximum Euclidean distance across all pairs of grid cells. Network analysis was then used to compute the shortest path across each grid cell’s climate-cost surface, producing a grid cell × grid cell matrix of least-cost distance, with values of each row representing the length of the shortest path of one focal grid cell to all other grid cells across its individual climate dissimilarity surface. The final distance between two grid cells was taken as the mean of the distances calculated when each grid cell was treated as focal. For more details on calculating least-cost distances, see Cai *et al*. (2025).

#### Trade flows

To quantify the effects of direct human activities driving species introductions between regions, we calculated bilateral trade flows (i.e., representing a suite of introduction pathways) between pairs of regions using data from the BACI international trade database (Gaulier & Zignago, 2010). BACI provides annual, product-level bilateral trade values (USD) for over 200 countries, classified under the Harmonized System (HS), the global standard for trade nomenclature. Although BACI covers the period from 1995 to 2023, we used 2022 data reported under the HS 2017 classification to ensure both data accuracy and a better global coverage. Each BACI record represents the trade value (USD) from one exporting country to one importing country for a specific product category. To obtain a single, undirected measure of bilateral trade flow for each country pair, we summed the trade values in both directions—that is, from country i to country j and from j to i—across all product categories. If data were available in only one direction, we used that value as the bilateral trade flow. We then matched regions in BACI to our regions, resulting in a dataset of 8,417 country pairs among 150 countries. To estimate trade flows between subregions located in different countries, which are below the country level and not represented in BACI, we weighted the national-level trade flow by the product of the population proportions of the two subregions, calculated as each subregion’s population divided by the total population of its respective country. The population was extracted as the total values per geographic region from the 2020 population count raster data (Center For International Earth Science Information Network-CIESIN-Columbia University, 2018).

### Identification of biogeographical realms

We conducted hierarchical clustering analyses of the compositional dissimilarity matrices to identify the floristic realms and subrealms globally, using the “hclust” function in the R package of stats. This method starts with each geographic region as an individual cluster, progressively merging clusters based on their compositional similarity until all regions belong to a single cluster. We used an unweighted pair group method with arithmetic mean (UPGMA), which has been shown to perform best among different clustering methods for quantitatively identifying biogeographical regionalization (Kreft & Jetz, 2010).

We applied the UPGMA method to taxonomic and phylogenetic turnover across two distinct datasets: native species and all species, generating four dendrograms. First, we identified native floristic realms and subrealms by cutting the dendrogram of native species at different heights based on a predetermined number of clusters. The number of clusters for native species datasets was determined by the within-between ratio, calculated as the ratio of the average within-cluster distance to the average between-cluster distance. Lower values of this ratio indicate better clustering performance. Specifically, we selected the number of clusters at which the within-between ratio showed a substantial decrease compared to the previous cluster number, followed by a relatively smaller change relative to the subsequent one (Fig. S12). This resulted in the identification of seven realms and 17 subrealms for taxonomic turnover, and eight realms and 18 subrealms for phylogenetic turnover (Fig. S12). Cluster names were assigned following Takhtajan’s scheme (Takhtajan, 1986) and established biogeographical terminology. However, the boundaries of our delineated realms and subrealms may differ from those of established schemes, as our analysis is based on species checklists compiled largely within political rather than strictly geographic borders. In addition, we evaluated clustering performance by calculating the proportion of beta diversity explained by the resulting clusters as the ratio between the sum of dissimilarities among regions of different clusters and the sum of all dissimilarities of all regions (Holt *et al*., 2013). For both taxonomic and phylogenetic turnover, floristic realms accounted for over 85% of the variation, whereas subrealms explained more than 93%. To explore whether and how the biogeographical patterns change due to non-native species, we then pruned the dendrogram of the all-species dataset at the dendrogram heights where native floristic realms and subrealms were delineated to make the patterns comparable.

To assess whether the number of biogeographical regions influences the detection of biogeographical pattern change due to plant introduction, we performed an additional clustering analysis using the same number of floristic realms (six) and subrealms (12) as defined in Takhtajan’s work (Takhtajan, 1986), across the native and all-species datasets (Figs. S13 and S14). Consistent with our main findings, these analyses also revealed reductions due to species introductions in the number of realms and subrealms in both taxonomic and phylogenetic compositions (Figs. S13 and S14). To assess whether correcting GIFT native species distribution based on continental-level native range from WCVP influenced the observed patterns, we also performed clustering analyses using the dataset without applying these corrections. The results were identical to those based on the dataset with WCVP native range corrections for taxonomic turnover at the realm level (Fig. S10 and Fig. 1), while phylogenetic turnover showed some differences at the realm and subrealm levels, though overall patterns remained consistent (Fig. S11 and Fig.2). These findings reinforce the robustness of the breakdown of biogeographical boundaries identified despite using partially different data.

### Individual non-native species contribution to variation in turnover

To assess the impact of individual non-native species on compositional dissimilarity and shifts in biogeographical patterns at the subrealm level, we quantified variation in taxonomic turnover due to the removal of each focal non-native species for each pair of subrealms. We did not perform this analysis at the realm level because of its relatively broad spatial scale. For each non-native species, we first removed its non-native distribution records from the dataset containing all species distributions. Using this modified dataset, we then calculated taxonomic turnover for each pair of geographic regions and computed the difference between these turnover values and those derived from the original dataset, where positive values indicated increased similarity driven by the presence of the focal species. To quantify the contribution of the focal species to variation in turnover between two natural subrealms (i.e., subrealms identified based on native species distribution), we averaged these turnover differences across all pairs of geographic regions belonging to the focal first and second subrealms, respectively. This procedure was repeated for each non-native species and for all subrealm pairs to estimate each species’ unique contribution to compositional homogenization between two specific subrealms. Higher values indicate greater similarity between subrealms resulting from the presence of the focal non-native species, reflecting a more pronounced impact on biogeographical patterns (Table S1). We then visualized the distribution patterns of two species with high contributions to the breakdown of subrealms, along with the regionalization patterns driven by their influence (Fig. S3).

### Null models

To assess the influence of potentially incomplete non-native species data—particularly in tropical regions—on biogeographical patterns found in this study, we implemented two alternative null model approaches based on taxonomic turnover. First, we tested whether the observed patterns would change if regions with low non-native species richness in our data harbored more non-native species, while all other regions remained unchanged. For regions with fewer than 100 non-native species, we randomly added 100 non-native species drawn from the non-native species pool of neighboring regions. These thresholds were chosen to 1) capture most potentially under-sampled regions in our data set, particularly in tropical and subtropical areas and 2) to add an amount of theoretically not yet detected but still possible to occur non-native species in those regions (Fig. S15a). Although the defined species pool here does not account for species-specific environmental requirements, we found that in most regions, over 80% of non-native species were also non-native in at least one of the ten geographically nearest regions (Fig. S16). To ensure the added species represented true non-local non-natives, we restricted the species pool to those non-native in at least one of the ten geographically nearest regions but native to none of them. We then recalculated taxonomic turnover and performed clustering analyses for the modified dataset that included both native and non-native species. The procedure was repeated 1000 times, resulting in 1000 distinct clustering outcomes to assess variability and robustness. (Fig. S4). Second, we tested whether biogeographical patterns change when the number of non-native species is reduced across regions, to assess the potential dominating effects of regions with intense sampling effort. For each region with more than 80 non-native plant species, we randomly removed 50 non-native species. These thresholds were chosen because the majority of regions were included based on the threshold (80% contained ≥80 non-native species; Fig. S15b), while allowing a substantial number of non-native species to be removed, sufficient to meaningfully affect regional composition and patterns. Clustering analyses were then performed on the modified dataset, and the procedure was repeated 1,000 times. This allowed us to assess whether the patterns remained consistent despite the reduced number of non-native species (Fig. S5).

### Statistical modeling

We used generalized dissimilarity modeling (GDM) to compare the contribution of dispersal limitation and environmental filtering in explaining spatial patterns of compositional dissimilarity (Ferrier *et al*., 2007). Specifically, we modelled taxonomic and phylogenetic turnover across different species groups for subsets of regions using different thresholds of minimum area sizes in response to environmental dissimilarity and geographical distances, and modelled additionally against trade flow. GDM is an effective tool for analyzing and predicting spatial patterns of beta diversity between pairs of regions, accounting for nonlinearities between explanatory variables and beta diversity (Mokany *et al*., 2022). The approach is capable of fitting (1) the curvilinear relationship between compositional turnover and increasing ecological and environmental distance between regions, and (2) the variation in the rate of compositional turnover at different positions along an environmental gradient (Ferrier *et al*., 2007). A fundamental assumption of GDM is that dissimilarity increases as the differences in predictor variable values between two regions become larger (Mokany *et al*., 2022).

#### The effects of dispersal limitation and environmental filtering

We fitted GDMs for taxonomic and phylogenetic turnover across three species distribution data (i.e., native, non-native, all-species datasets), respectively, using the R package *gdm* (Fitzpatrick *et al*., 2022) with the region × region matrices of geographical distances (i.e., geographical linear distances, least-cost distance across all barriers, and least-cost distance across climate dissimilarity surface), and the untransformed vectors of environmental variables (i.e., mean annual temperature, mean annual precipitation, temperature seasonality, precipitation seasonality, aridity index, and soil cation-exchange capacity) as predictors. We utilized three *I-spline* basis functions for each predictor. To test whether our results are affected by differences in area size among regions, we fitted GDMs for subsets of regions with minimum area sizes of 10, 1000, 10,000 and 100,000 km^2^, respectively (Fig.3, Fig. S6 and S7). Because patterns were consistent across subsets of regions with different area-size thresholds, we presented main results (Fig. 3) based on models including regions with a minimum area of 10 km².

Following König et al. (2017), we utilized two approaches to quantify the relative importance of predictors. First, we estimated the effect of each individual predictor in explaining turnover using the maximum height of fitted spline functions in the GDM (Fitzpatrick *et al*., 2022). We then scaled the heights such that their sums equaled the proportion of deviance explained by the model. Second, we used deviance partitioning to assess the relative importance of environmental filtering versus dispersal limitation (Borcard *et al*., 1992). Specifically, we partitioned the explained variation in turnover into four components: the fraction explained only by environmental filtering, the fraction explained only by dispersal limitation, the overlapping fraction explained by both dispersal limitation and environmental filtering, and the unexplained variation. This was achieved using the function *gdm.partition.deviance*.

#### The effects of trade flow

To assess the impact of trade flow, representing different introduction pathways, on compositional dissimilarity, we first fitted GDM models, additionally including bilateral trade flows. Because GDM cannot accommodate predictors with missing values, we assigned a trade value of zero to country pairs without BACI trade data, as well as to trade flows between subregions within the same country. Although zero represents the absence of trade rather than true missing data, this approach was necessary since GDM requires predictors to be structured in matrix form. Retaining zeros preserved a complete dataset of 115,921 region pairs (10,457 with zero values) across 482 regions, whereas excluding the 475 regions (86.5%) lacking complete trade flows would have drastically reduced coverage. In the model, we found that trade flow exerted a weak but detectable effect on taxonomic and phylogenetic turnover for all and non-native species (Fig. S8 and S9).

To avoid potential artifacts from our handling of missing trade data, we next tested the effects of bilateral trade flows on the difference in shared species between region pairs, calculated as the number of shared species after introduction minus that before introduction (i.e., values >= 0), using generalized linear models. This analysis was restricted to 8,417 region pairs covering 148 regions for which trade data were directly available in the BACI dataset, without rescaling to subregions or assigning zero values. In the model, we used a negative binomial error distribution with a log link function for difference in shared species to cope with the overdispersion of the response variable. Besides trade flow, we included contemporary climate differences between each pair of regions to capture the potential environmental limitations on the establishment of non-native species among regions. The climate difference of region pairs was defined as the Euclidean distance in a multidimensional climatic space, incorporating mean annual temperature, mean annual precipitation, temperature seasonality, precipitation seasonality, and aridity index. Before fitting the model, climatic distance and trade flow were log-transformed owing to their skewed distributions. After transformation, all continuous predictor variables were standardized to zero mean and unit variance to aid model fitting and make their parameter estimates comparable. To test whether the establishment of non-native species in a given region is influenced by the joint effects of climate and trade, we included an interaction term between climatic difference and trade flow in the model. However, based on AIC, the model including the interaction term performed worse than the model without it; therefore, we present results from the model excluding the interaction term.

## Acknowledgements

L.C., H.B., I.K., and M.W. acknowledge the German Research Foundation (DFG) funding via iDiv (DFG FZT 118, 202548816). F.E. and B.L. appreciate funding by the Austrian Science Fund FWF (pr. no.: I 5825-B). M.v.K and A.J.D. acknowledge the German Research Foundation (DFG; grant 264740629). PP and JP were supported by long-term research development project RVO 67985939 (Czech Academy of Sciences).

## Author contributions

L.C., P.W., H.K., and M.W. conceived the idea and developed the conceptual framework of the study. L.C., P.W., H.K., W.D., F.E., M.v.K., J.P., P.P., A.J.D., P.B.P., J.J.W., and M.W. compiled the data. L.C. and PW performed the analyses, with suggestions from H.B., I.K., and M.W.. L.C. and M.W. wrote the first draft of the manuscript. All authors contributed to the writing and interpretation of the results.

