## Supplementary Materials for "Global disruption of plant biogeography by non-native species"

Lirong Cai et al.

**The PDF file includes:**

Figs. S1 to S16

Table S1

References

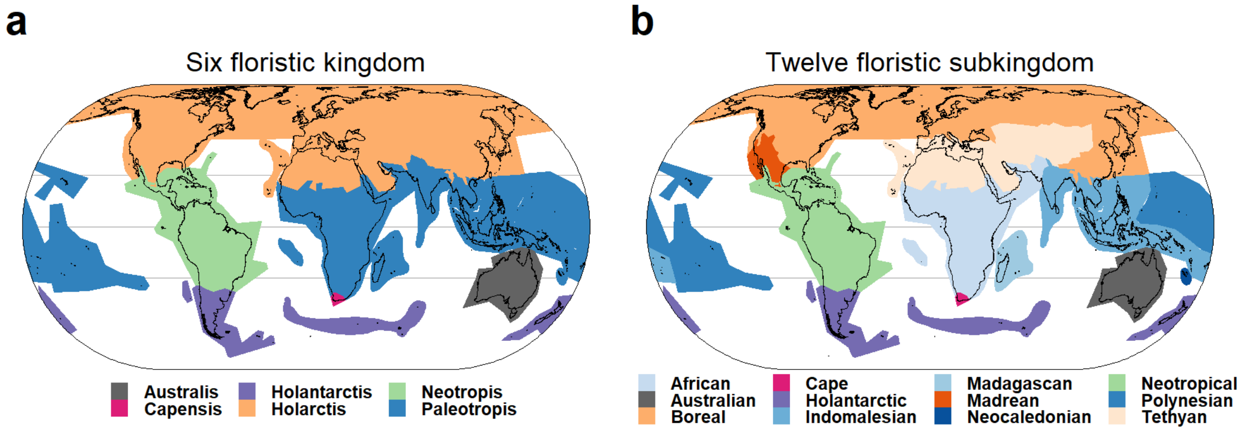


**Fig. S1. Takhtarjan's floristic scheme** (Takhtajan, 1986)**.** The global flora was divided into (a) six floristic kingdoms and (b) 12 subkingdoms.

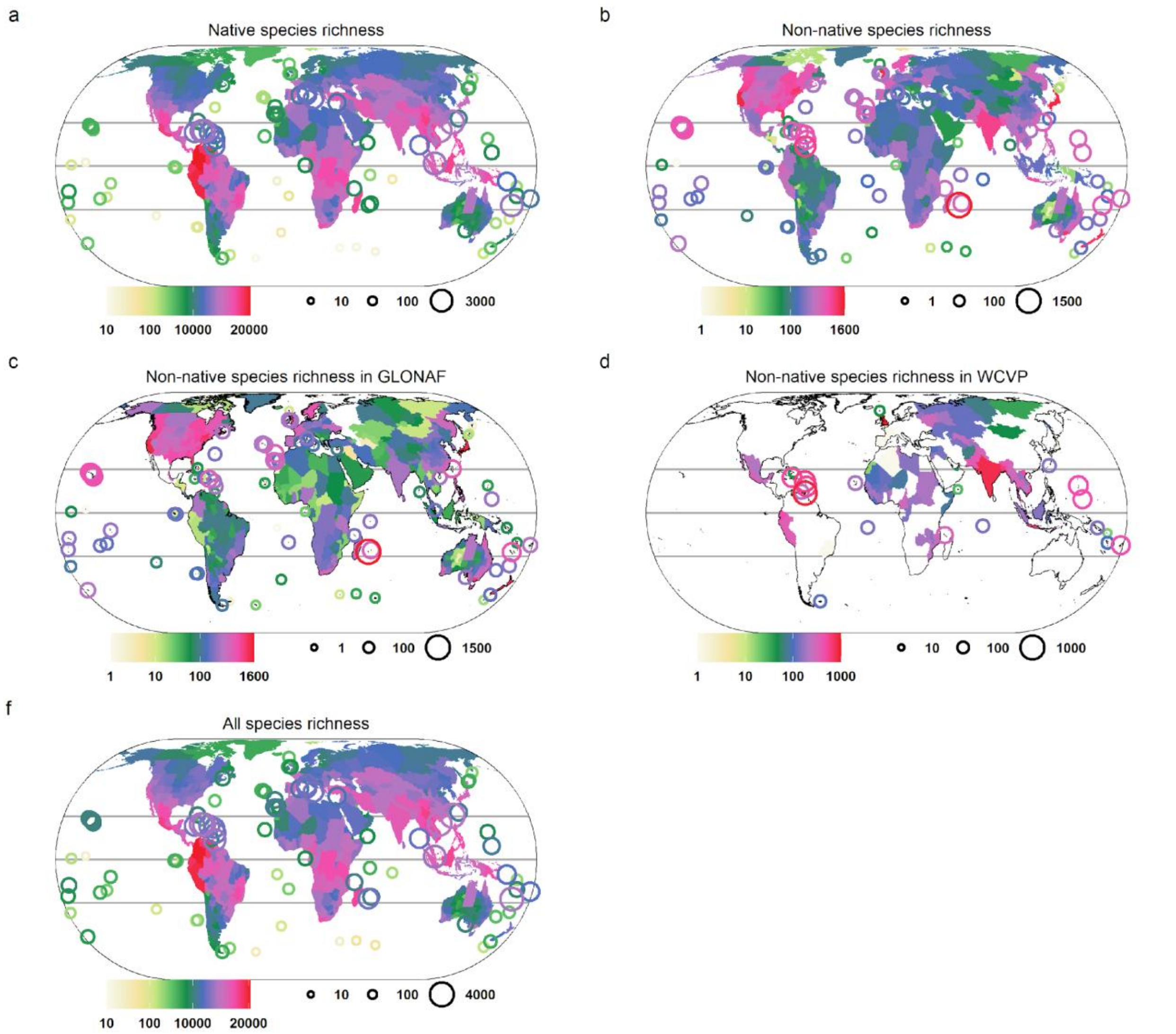


**Fig. S2. Observed species richness of seed plants across 549 geographic regions.** (a) Native species richness based on the Global Inventory of Floras and Traits (GIFT) (Weigelt *et al.*, 2020) and the World Checklist of Vascular Plants (WCVP) (Govaerts *et al.*, 2021). (b) Non-native species richness, compiled from (c) the Global Naturalized Alien Flora (GloNAF) (van Kleunen *et al.*, 2019) and (d) WCVP. (f) All species richness, combining native and non-native species. Species richness in panels (a–f) is shown on a $\log_{10}$ scale. All maps use the Eckert IV projection.

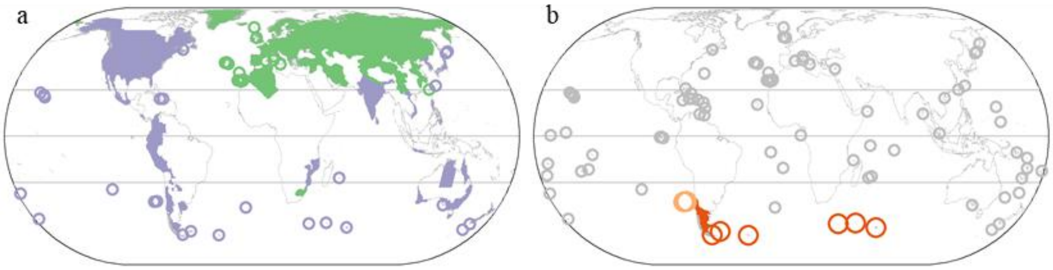


**Fig. S3. Example of a non-native plant species (*Rumex acetosella* L.) contributing to the breakdown of biogeographical boundaries.** (a) Native (green) and non-native (purple) ranges of the species. (b) The two subrealms (dark and light orange) where the species' non-native distribution contributed to boundary erosion.

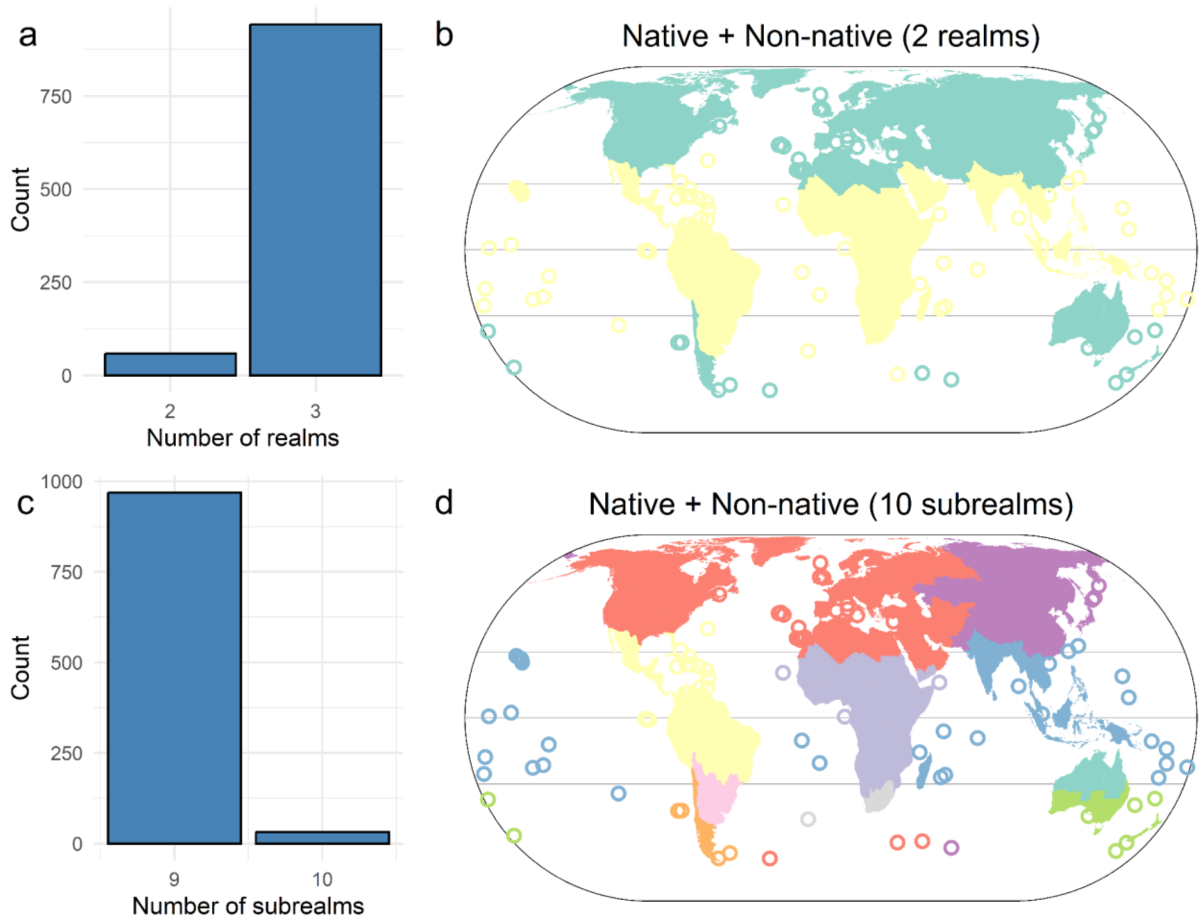


**Fig. S4. Results of the null model simulating species additions to focal regions from neighboring regions based on taxonomic turnover.** For regions with fewer than 100 non-native species, we randomly added 100 non-native species drawn from the non-native species pool of neighboring regions. Shown are the distributions of the number of biogeographical realms (a) and subrealms (c) after plant introduction across 1,000 iterations of the null model. Maps (b, d) illustrate the corresponding global biogeographical patterns of realms and subrealms for the runs that deviated from the actually observed patterns in Fig. 1.

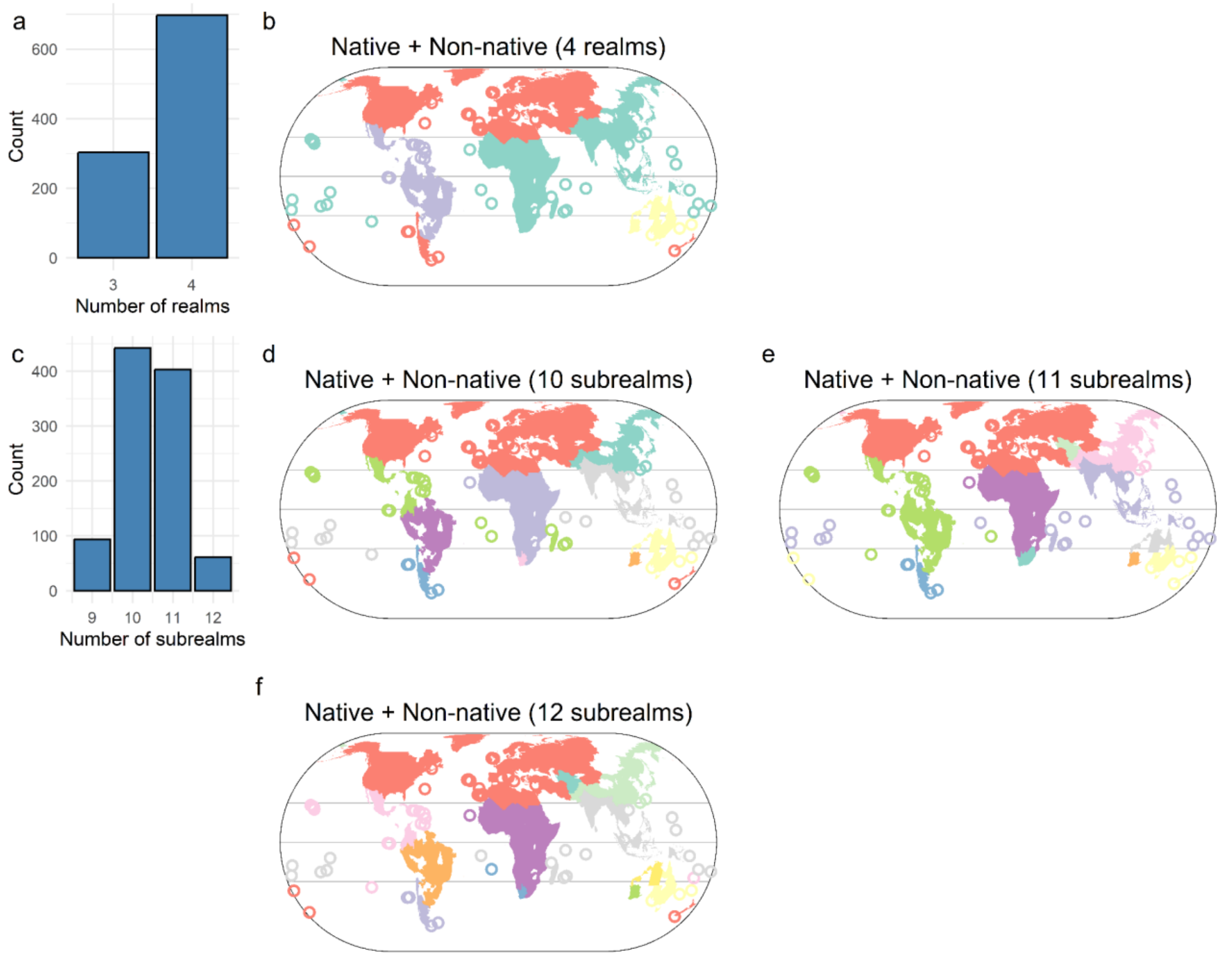


**Fig. S5. Results of the null model simulating species removal from focal regions based on taxonomic turnover.** For each region with more than 80 non-native plant species, we randomly removed 50 non-native species. Shown are the distributions of the number of biogeographical realms (a) and subrealms (c) after plant introduction across 1,000 iterations of the null model. Maps (b, d, e,f) illustrate the corresponding global biogeographical patterns of realms and subrealms for the runs that deviated from the actually observed patterns in Fig. 1.

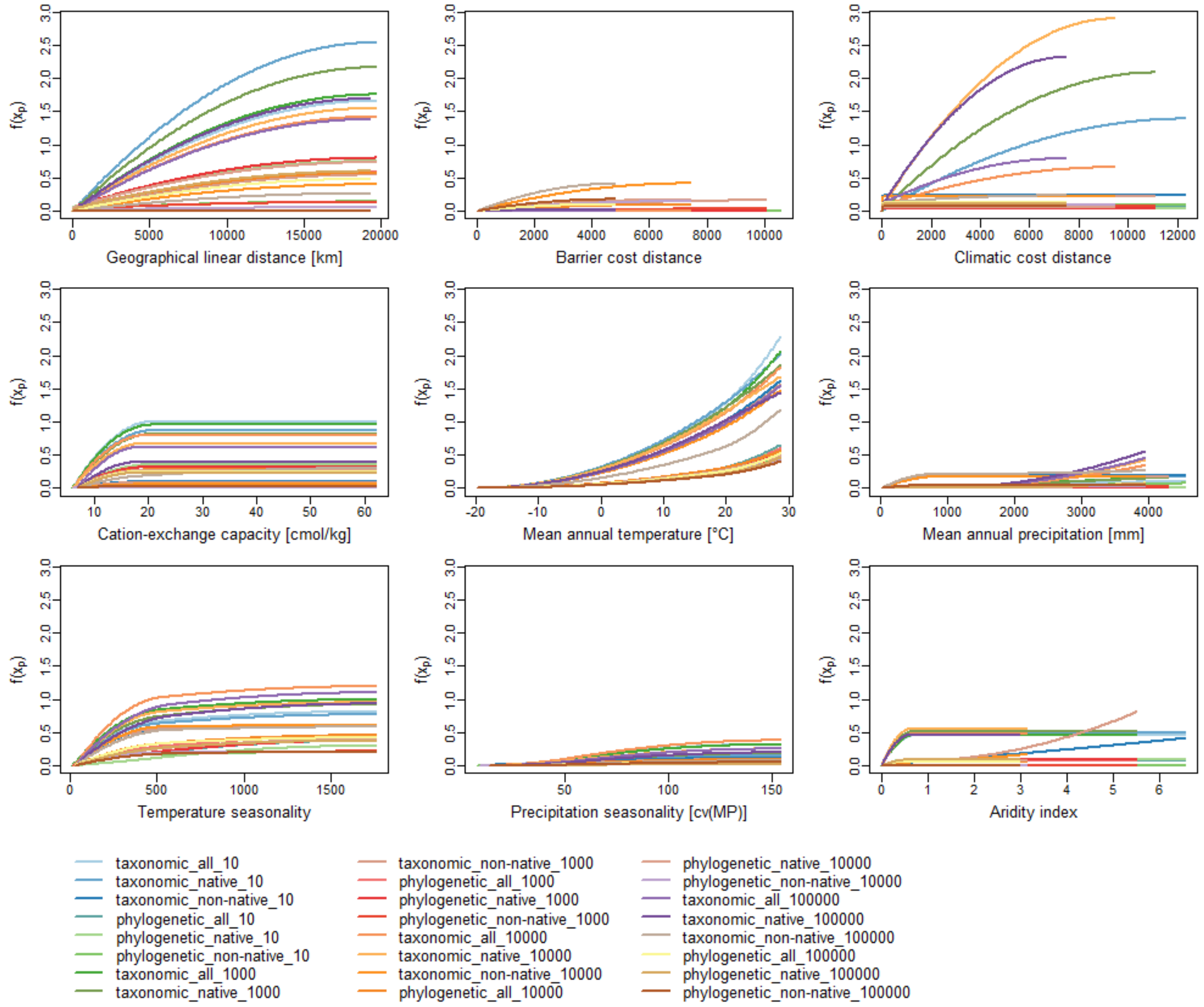


**Fig. S6. Spline functions from generalized dissimilarity modeling for each predictor variable for phylogenetic and taxonomic turnover in seed plants.** Analyses are carried out across native, non-native, and all species groups, and for regions with different minimum area sizes (10; 1,000; 10,000; 100,000 km²). The maximum height of the spline function indicates the importance of the predictor variable for explaining dissimilarities.

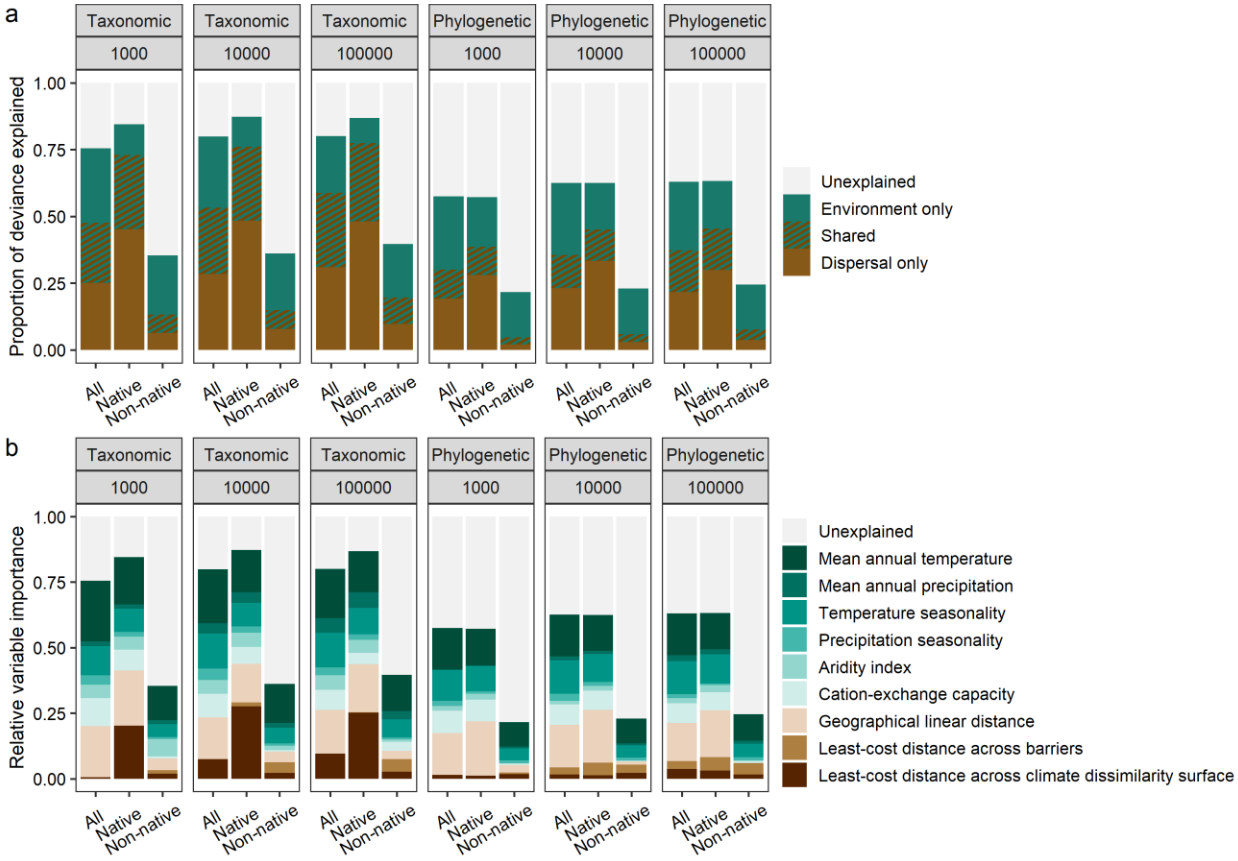


**Fig. S7. Relative importance of predictor variables for phylogenetic and taxonomic turnover in seed plants based on generalized dissimilarity models including regions with different minimum area sizes (i.e., 1,000; 10,000; 100,000 km$^2$).** In (a), relative importance is shown for environment and dispersal-related groups of predictor variables based on deviance partitioning. In (b), relative importance is shown for individual predictor variables based on the height of generalized dissimilarity modeling transformation curves, scaled to make their sums equal to the proportion of deviance explained by the model.

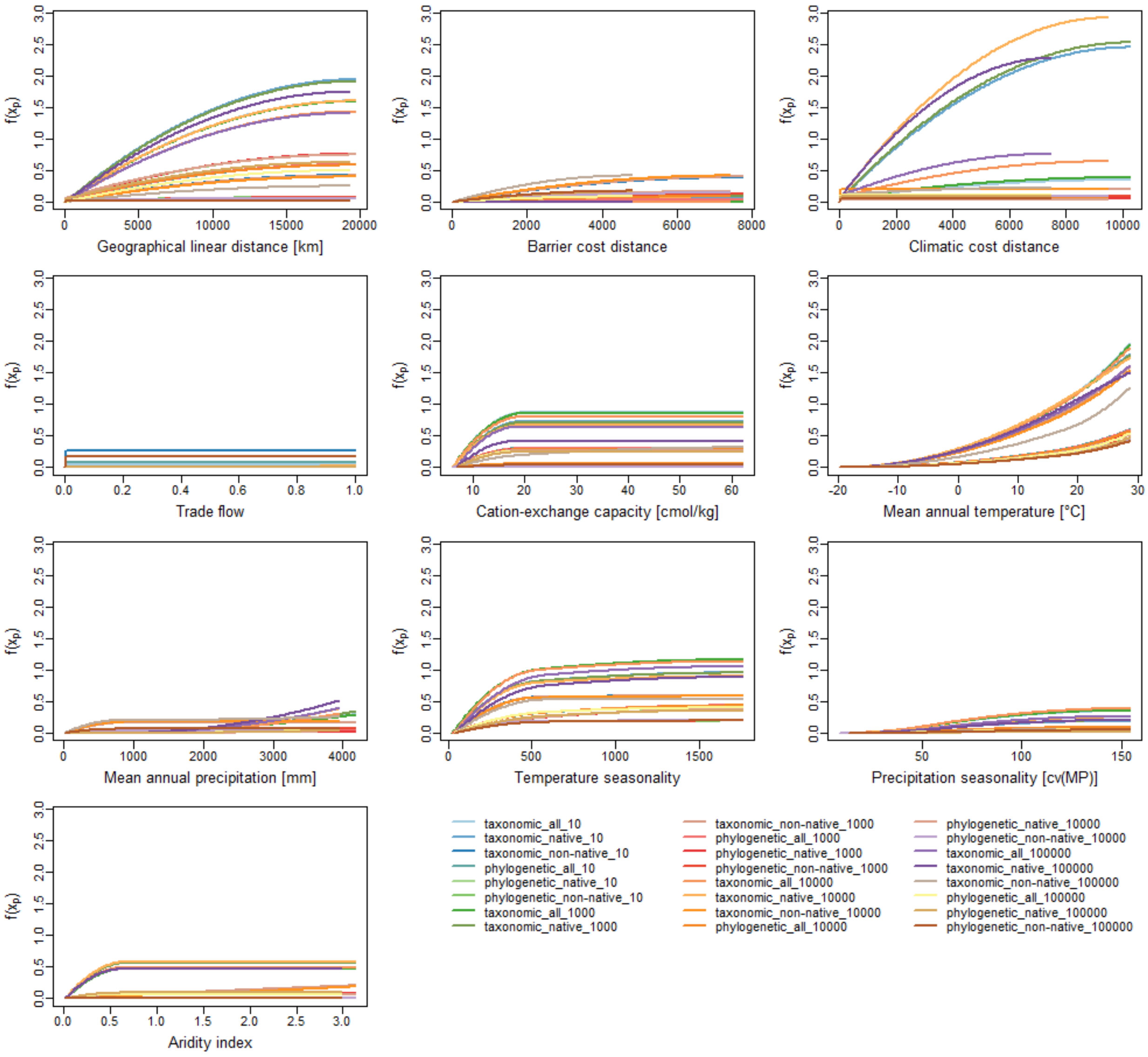


**Fig. S8. Spline functions from generalized dissimilarity modeling with bilateral trade flow for each predictor variable for phylogenetic and taxonomic turnover in seed plants.** Analyses are carried out across native, non-native, and all species groups, and for regions with different minimum area sizes (10; 1,000; 10,000; 100,000 km²). The maximum height of the spline function indicates the importance of the predictor variable for explaining dissimilarities.

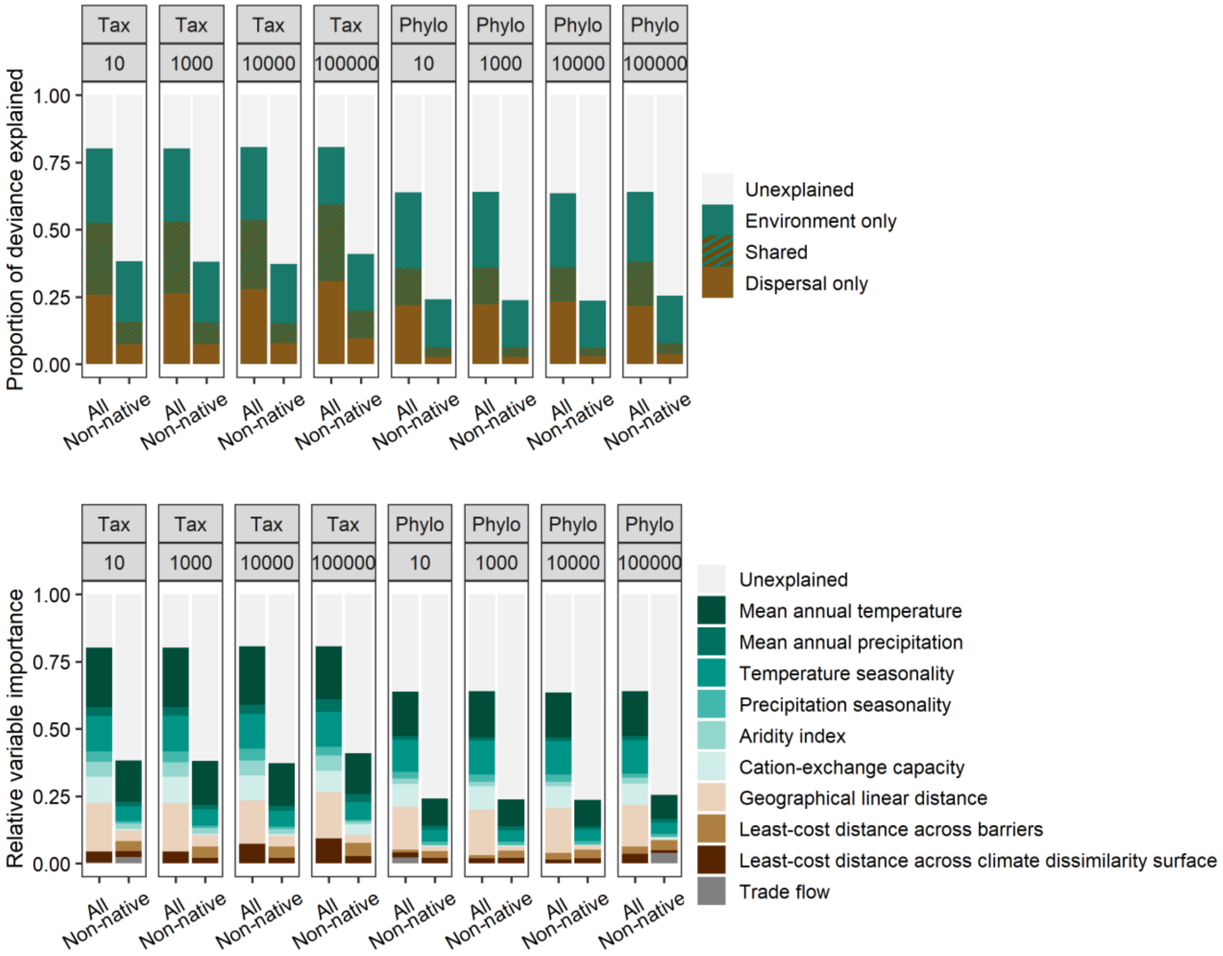


**Fig. S9. Relative importance of predictor variables for phylogenetic (Phylo) and taxonomic (Tax) turnover in seed plants based on generalized dissimilarity models with trade flow.** In (a), relative importance is shown for environment and dispersal-related groups of predictor variables based on deviance partitioning. In (b), relative importance is shown for individual predictor variables based on the height of generalized dissimilarity modeling transformation curves, scaled to make their sums equal to the proportion of deviance explained by the model. Variable importance is shown for models including regions with different minimum area sizes (i.e., 10; 1,000; 10,000; 100,000 km$^2$).

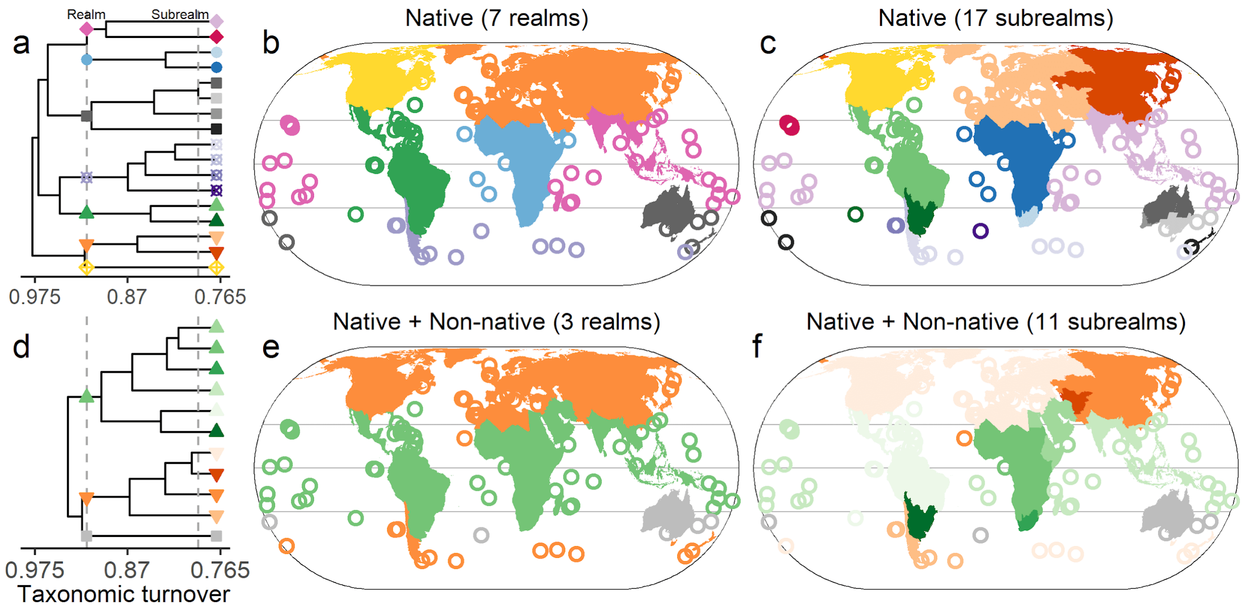


**Fig. S10. Global biogeographical patterns before and after plant introductions, based on taxonomic turnover derived from species distribution data without applying native range corrections from WCVP to the GIFT dataset.** Compositional dissimilarity was calculated using two species datasets: native species only (a–c) and all species combined (d–f). Dendrograms from hierarchical clustering (a, d) were pruned at two different heights to define biogeographical realms and subrealms. Realms are shown as distinct colors on the maps (b, e), while subrealms belonging to the same realm are depicted using consistent color gradients from the same palette on the maps (c, f) and are represented by the same symbols on the dendrograms. Matching colors are used to indicate corresponding realms and subrealms across the dendrograms and maps. Islands <50,000 km$^2$ are represented by circles.

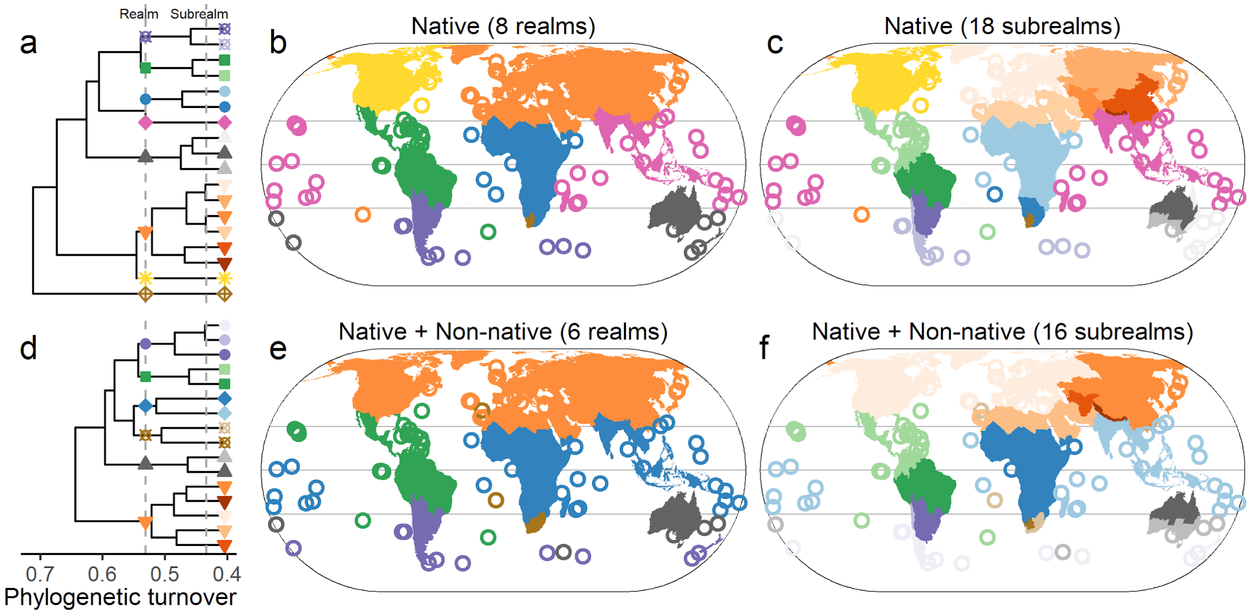


**Fig. S11. Global biogeographical patterns before and after plant introductions, based on phylogenetic turnover derived from species distribution data without applying native range corrections from WCVP to the GIFT dataset.** Compositional dissimilarity was calculated using two species datasets: native species only (a–c) and all species combined (d–f). Dendrograms from hierarchical clustering (a, d) were pruned at two different heights to define biogeographical realms and subrealms. Realms are shown as distinct colors on the maps (b, e), while subrealms belonging to the same realm are depicted using consistent color gradients from the same palette on the maps (c, f) and are represented by the same symbols on the dendrograms. Matching colors are used to indicate corresponding realms and subrealms across the dendrograms and maps. Islands <50,000 km$^2$ are represented by circles.

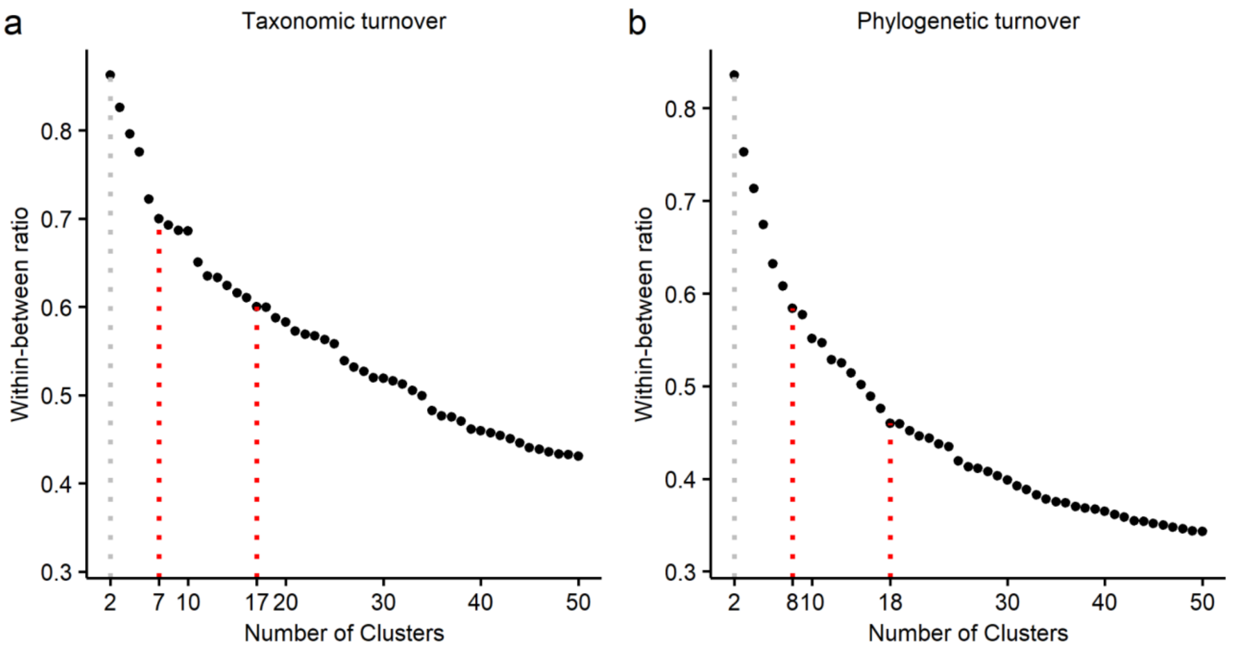


**Fig. S12. Within-between ratio plotted against the number of clusters for (a) taxonomic turnover and (b) phylogenetic turnover.** Each data point represents the ratio corresponding to a given number of clusters. Red dotted vertical lines indicate selected cluster numbers based on their associated within–between ratios.

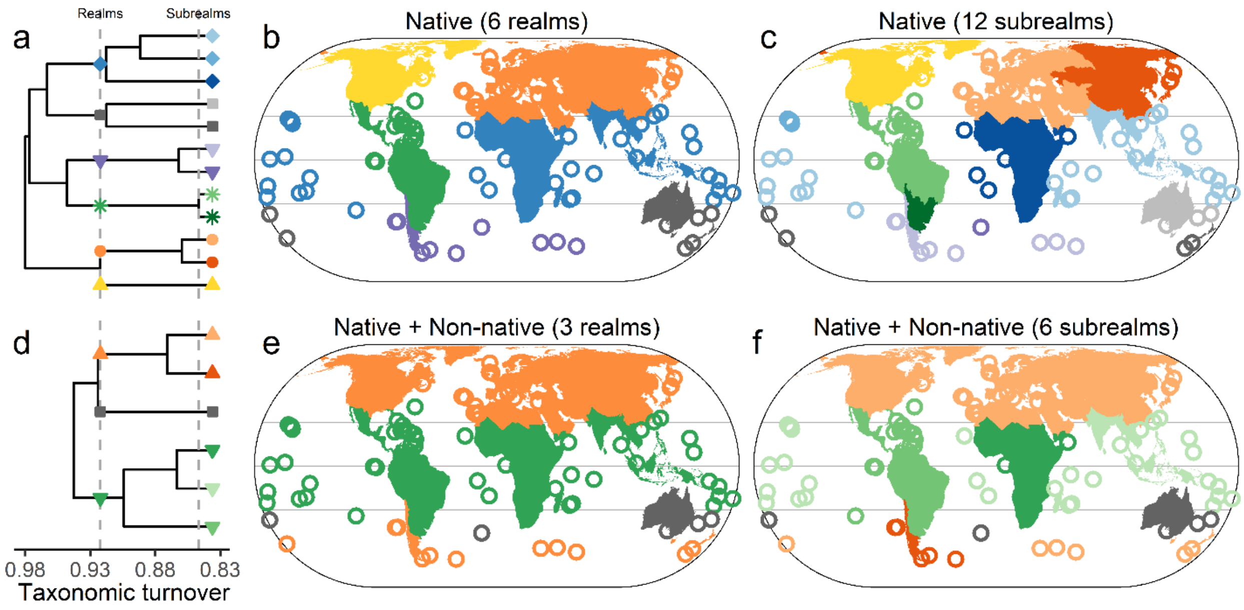


**Fig. S13. Global biogeographical patterns before and after plant introduction based on taxonomic turnover. The number of clusters for native species was determined according to Takhtajan's floristic scheme** (Takhtajan, 1986)**.** Compositional dissimilarity was calculated using two species datasets: native species only (a–c) and all species combined (d–f). Dendrograms from hierarchical clustering (a, d) were pruned at two different heights to define biogeographical realms and subrealms. Realms are shown as distinct colors on the maps (b, e), while subrealms belonging to the same realm are depicted using consistent color gradients from the same palette on the maps (c, f) and are represented by the same symbols on the dendrograms. Matching colors are used to indicate corresponding realms and subrealms across the dendrograms and maps. Islands <50,000 km$^2$ are represented by circles.

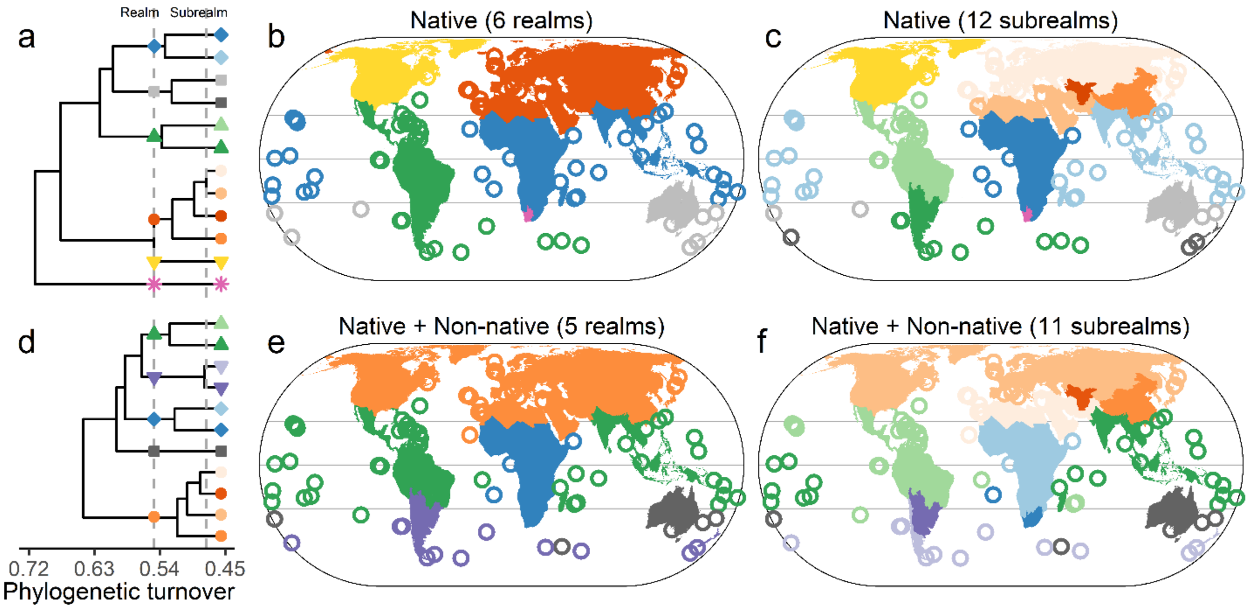


**Fig. S14. Global biogeographical patterns before and after plant introduction based on phylogenetic turnover. The number of clusters for native species was determined according to Takhtajan's floristic scheme** (Takhtajan, 1986)**.** Compositional dissimilarity was calculated using two species datasets: native species only (a–c) and all species combined (d–f). Dendrograms from hierarchical clustering (a, d) were pruned at two different heights to define biogeographical realms and subrealms. Realms are shown as distinct colors on the maps (b, e), while subrealms belonging to the same realm are depicted using consistent color gradients from the same palette on the maps (c, f) and are represented by the same symbols on the dendrograms. Matching colors are used to indicate corresponding realms and subrealms across the dendrograms and maps. Islands <50,000 $km^2$ are represented by circles.

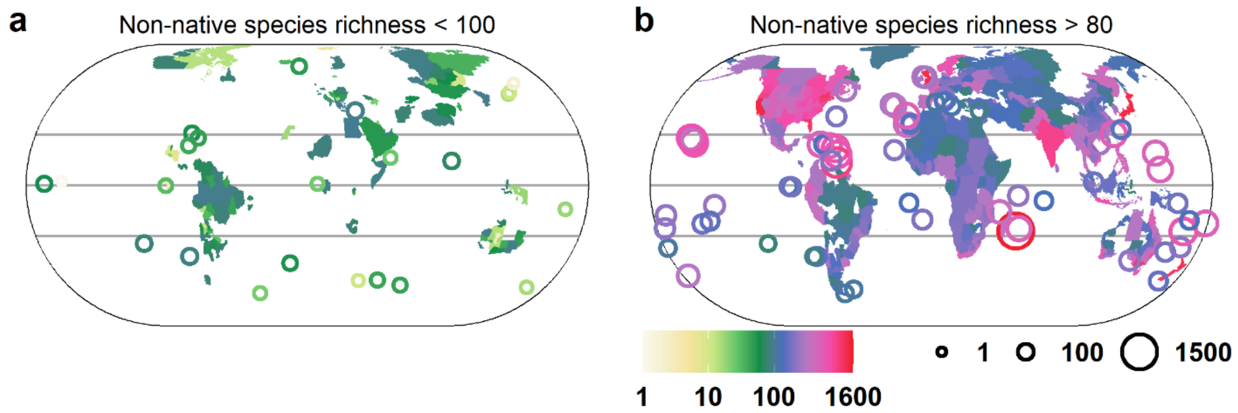


**Fig. S15. Observed regions (a) with fewer than 100 non-native seed plant species and (b) with more than 80 non-native seed plant species.**

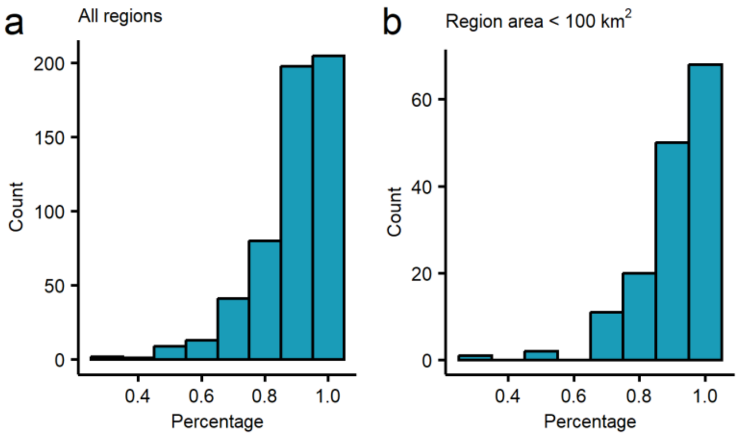


**Fig. S16. The frequency distribution of the percentage of non-native species in the focal region that are non-native to at least one neighboring region.** Neighboring regions for each focal region are defined as the ten closest regions based on great-circle distance.

**Table S1. The top five species contributing to the breakdown of biogeographical boundaries for different subrealm pairs.**

| First subrealm | Second subrealm | Species |
|---|---|---|
| Eastern Australia | Tristan da Cunha | *Sonchus oleraceus* |
| | | *Cynodon dactylon* |
| | | *Chenopodiastrum murale* |
| | | *Lysimachia arvensis* |
| | | *Dysphania ambrosioides* |
| Eastern Australia | Western Australia | *Sonchus oleraceus* |
| | | *Cynodon dactylon* |
| | | *Tribulus terrestris* |
| | | *Citrullus lanatus* |
| | | *Heliotropium curassavicum* |
| Eastern Australia | Southwest Australia | *Sonchus oleraceus* |
| | | *Chenopodiastrum murale* |
| | | *Centaurea melitensis* |
| | | *Echium plantagineum* |
| | | *Lysimachia arvensis* |
| Europe | New Zealand | *Sonchus oleraceus* |
| | | *Stellaria media* |
| | | *Rumex acetosella* |
| | | *Cerastium glomeratum* |
| | | *Juncus bufonius* |
| Europe | North America | *Capsella bursa-pastoris* |

| | | |
|---|---|---|
| | | *Stellaria media* |
| | | *Rumex crispus* |
| | | *Raphanus raphanistrum* |
| | | *Sonchus asper* |
| Hawaii | South Asia | *Heliotropium arboreum* |
| | | *Cyperus brevifolius* |
| | | *Eleusine indica* |
| | | *Cyanthillium cinereum* |
| | | *Cyperus mindorensis* |
| New Zealand | North America | *Stellaria media* |
| | | *Cerastium fontanum* |
| | | *Rumex acetosella* |
| | | *Juncus bufonius* |
| | | *Sonchus oleraceus* |
| South South America | Juan Fernádez | *Rumex acetosella* |
| | | *Plantago lanceolata* |
| | | *Agrostis stolonifera* |
| | | *Lolium multiflorum* |
| | | *Stellaria media* |
| South South America | North Chile | *Agrostis stolonifera* |
| | | *Plantago lanceolata* |
| | | *Medicago sativa* |
| | | *Rumex acetosella* |

| | | |
|---|---|---|
| | | *Lobelia oligophylla* |
| Tristan da Cunha | Western Australia | *Sonchus oleraceus* |
| | | *Cynodon dactylon* |
| | | *Pseudognaphalium luteoalbum* |
| | | *Lepidium didymum* |
| | | *Chenopodiastrum murale* |
| Tristan da Cunha | Southwest Australia | *Sonchus oleraceus* |
| | | *Chenopodiastrum murale* |
| | | *Malva parviflora* |
| | | *Lysimachia arvensis* |
| | | *Juncus bufonius* |
| Juan Fernández | North Chile | *Galinsoga parviflora* |
| | | *Sonchus oleraceus* |
| | | *Chenopodiastrum murale* |
| | | *Medicago polymorpha* |
| | | *Briza minor* |
| Western Australia | Southwest Australia | *Sonchus oleraceus* |
| | | *Tribulus terrestris* |
| | | *Salsola tragus* |
| | | *Cynodon dactylon* |
| | | *Sisymbrium orientale* |
